# Nod1-dependent NF-kB activation initiates hematopoietic stem cell specification in response to small Rho GTPases

**DOI:** 10.1101/2022.11.19.517153

**Authors:** Xiaoyi Cheng, Radwa Barakat, Abigail Gorden, Elizabeth Snella, Yudi Zhang, Karin Dorman, Antonella Fidanza, Clyde Campbell, Raquel Espin-Palazon

## Abstract

The possibility of specifying functional hematopoietic stem and progenitor cells (HSPCs) from human pluripotent stem cells (hPSCs) would overcome current limitations related to HSPC transplantation. However, generating hPSC-derived HSPCs has been elusive, necessitating a better understanding of the native developmental mechanisms that trigger HSPC specification. Here, we revealed *in vivo* an intrinsic inflammatory mechanism triggered by Nod1 that drives early hemogenic endothelium (HE) patterning to specify HSPCs. Our genetic and chemical experiments showed that HSPCs failed to specify in the absence of Nod1 and its downstream kinase Ripk2. Rescue experiments demonstrated that Nod1 and Ripk2 acted through NF-kB, and that small Rho GTPases are at the apex of this mechanism. Manipulation of NOD1 in a human system of hPSCs differentiation towards the definitive hematopoietic lineage indicated functional conservation. This work establishes the RAC1-NOD1-RIPK2-NFkB axis as the earliest inflammatory inductor that intrinsically primes the HE for proper HSPC specification. Manipulation of this pathway could help derive a competent HE amenable to specify functional patient specific HSPCs for the treatment of blood disorders.

**Highlights:**

1. Nod1 specifies HSPCs *in vivo* through the early induction of hemogenic endothelium.
2. Nod1-Ripk2 controls HSPC specification by activating the inflammatory master TF NF-kB.
3. Nod1 links small Rho GTPases with pro-inflammatory signaling during the genesis of HSPCs.
4. The function of NOD1 is conserved in the development of definitive human HSPCs.

## Introduction

In the context of injury and infection, Pattern Recognition Receptors (PRRs) sense Pathogen-Associated Molecular Patterns (PAMPs) to activate Nuclear Factor-kB (NF-kB) and trigger the expression of pro-inflammatory cytokines. These in turn will activate the immune system to eliminate the infection and preserve the integrity of the organism. However, in the last few years it has become clear that the vertebrate embryo, in the absence of injury or infection, also utilizes pro-inflammatory signaling as a developmental mechanism to specify hematopoietic stem and progenitor cells (HSPCs) (Espin-Palazon et al., 2014; Frame et al., 2020; He et al., 2015; Lefkopoulos et al., 2020; Li et al., 2014; Sawamiphak et al., 2014; Tie et al., 2019), a phenomenon referred to as developmental inflammation (Espin-Palazon et al., 2018). Although HSPCs and myeloid cell are an important source of pro-inflammatory cytokines in this context (Espin-Palazon et al., 2014; Li et al., 2014; Tie et al., 2019), the drivers of this “embryonic cytokine storm” are largely unknown. Identifying how and when developmental inflammation is initiated in the embryo to generate HSPCs is a critical need to achieve the elusive goal of generating functional HSPCs from human pluripotent stem cells (hPSCs) for therapeutic use.

The genesis of HSPCs from hemogenic endothelium (HE) is conserved across vertebrate phyla (Ivanovs et al., 2017). During a brief window of embryonic development, HSPCs arise *de novo* from hemogenic endothelium (HE) comprising the ventral floor of the dorsal aorta (vDA), a process termed the endothelial to hematopoietic transition (EHT). During EHT, endothelial cells undergo morphological and genetic changes that give rise to HSPCs (Bertrand et al., 2010; Boisset et al., 2010; de Bruijn et al., 2000; Kissa and Herbomel, 2010; Zovein et al., 2008). Current human differentiation protocols have been largely extrapolated from animal models, including zebrafish (Frame et al., 2017; Rowe et al., 2016), and many clinical trials have been derived from findings in zebrafish (Cully, 2019; North et al., 2007). In addition, zebrafish embryos develop externally, circumventing the artifactual inflammatory conditions triggered by surgical procedures needed in mammalian models, and providing an ideal model to investigate inflammatory signaling during embryogenesis.

PRRs are proteins classically known for their ability to recognize pathogen signatures, also known as PAMPs, and trigger the expression and release of pro-inflammatory cytokines to activate the immune system to fight the insult. Among them, the NOD-Leucin Rich Repeats (LRR)-containing receptors (NLRs) are one of the major sub-family of PRRs. NLRs are highly conserved cytosolic PRRs that scrutinize the intracellular environment for the presence of infectious or harmful perturbations. Once sensed, NLRs oligomerize, forming macromolecular scaffolds to recruit effector signaling cascades that lead to inflammation (Geddes et al., 2009; Ting et al., 2008). Non-inflammasome forming NLRs, which include NOD1, NOD2, NLRP10, NLRX1, NLRC5, and CIITA, do not engage the inflammasome, but rather activate NF-kB, mitogen-activated protein kinases (MAPKs), and interferon regulatory factors (IRFs) (Zhong et al., 2013). Classically, NOD1 is activated by diaminopimelic acid (DAP)-type peptidoglycan, a type of bacterial peptidoglycan found almost exclusively in Gram-negative bacteria. Once activated, Nod1 oligomerizes and recruits RIPK2 leading to activated NF-kB and MAPK signaling that leads to classical inflammation (Chamaillard et al., 2003; Kobayashi et al., 2002). In addition, it has been demonstrated recently that NOD1 and NOD2 regulate adult stem cell function (Burberry et al., 2014; Nigro et al., 2014). Specifically, NOD1 can mobilize adult HSPCs in a murine model of bacteremia (Burberry et al., 2014). Whether or not non-inflammasome forming NLRs, and particularly NOD1, induce developmental inflammation to specify HSPCs in the aseptic embryo has not been investigated. To address this need, we have taken advantage of the external embryonic development of the zebrafish and identified *in vivo* an inflammatory mechanism that serves as an inductive signal to make competent HE that will produce HSPCs *de novo*. Unexpectedly, we found that NF-kB, the central hub of pro-inflammatory signaling, was induced during HE patterning, a much earlier time than previously captured. We demonstrated that zebrafish and human HE expressed *NOD1* and its downstream effectors, providing a plausible candidate pathway that could power NF-kB activation during the patterning of HE. By performing genetic and chemical ablation of Nod1 and Ripk2, we found that this inflammatory mechanism was indeed crucial for early HE patterning prior to EHT. Rescue experiments demonstrated that small Rho GTPases induced the Nod1-Ripk2-NF-kB inflammatory mechanism in the HE prior blood flow. Last, we showed that NOD1 activation was also required to generate human definitive hematopoietic progenitors, highlighting a clear conservation for this mechanism across phyla. In summary, we identified the molecular mechanism by which developmental inflammation is first initiated in the embryo to generate HSPCs, as well as assigned a previously unappreciated function for inflammatory signaling during early HE patterning.

## Results

### Early HE activates NF-kB and expresses non-inflammasome-forming NLRs

HSPC development is a complex and highly dynamic process occurring only during embryonic development that involves the following developmental trajectory: (1) mesoderm formation; (2) endothelial and artery fate determination; (3) HE induction; (4) endothelial to hematopoietic (EHT) transition; and (5) HSPC release from the vDA through epithelial to mesenchymal transition (EMT) (Fig. 1A) (REFs). In previous studies, we utilized an NF-kB reporter zebrafish line, *Tg(NF-kB:eGFP)* (Kanther et al., 2011), in combination with the *kdrl:mcherry* transgene (Bertrand et al., 2010) to monitor NF-kB activation in the DA. We demonstrated that NF-kB was active in the DA during EHT (24 and 30 hpf) (Espin-Palazon et al., 2014). However, the temporal initiation of NF-kB signaling was unclear. To address this, we live imaged the HE by confocal imaging during its early (16 hours post fertilization, hpf) and late induction (20 hpf) using *Tg(kdrl:mcherry; NF-kB:eGFP)* zebrafish embryos. As shown in Fig. 1B, NF-kB activation in the HE was observed modestly at 16 hpf, and increased overtime (Fig. 1C), demonstrating that NF-kB signaling activated during HE induction previous EHT. We reasoned that, because primitive myeloid cells are not found within the DA boundary this early during development, an intrinsic inflammatory mechanism must drive NF-kB activation endogenously within the HE. Since non-inflammasome forming NLRs are one of the main NF-kB inductors during classical inflammation, we queried if they were expressed by the HE during its induction. All non-inflammasome-forming NLRs transcripts, except for *nlrc5*, were enriched in fluorescence activated cell sorted (FACS) purified *kdrl*^*+*^ HE from 22 (hpf) *kdrl:mcherry* zebrafish embryos as assessed by qPCR (Fig. 1D). Among them, *NOD1, CIITA*, and *NLRX1* were indeed expressed in human vascular HE cells at Carnegie stage (CS) 12-14 by scRNA-seq from a previously published dataset (Zeng et al., 2019), although *NOD1* transcripts were the most abundant among all (Fig. 1E-H). Interestingly, *RIPK2*, the main downstream kinase effector of NOD1, and *NFKBIA*, a surrogate of NF-kB activation, were also highly expressed (Fig 1I-J). These data suggested a potential and novel role for non-inflammasome forming NLRs during early HE induction, and particularly for NOD1 and its downstream signaling pathway.

**Figure 1.**
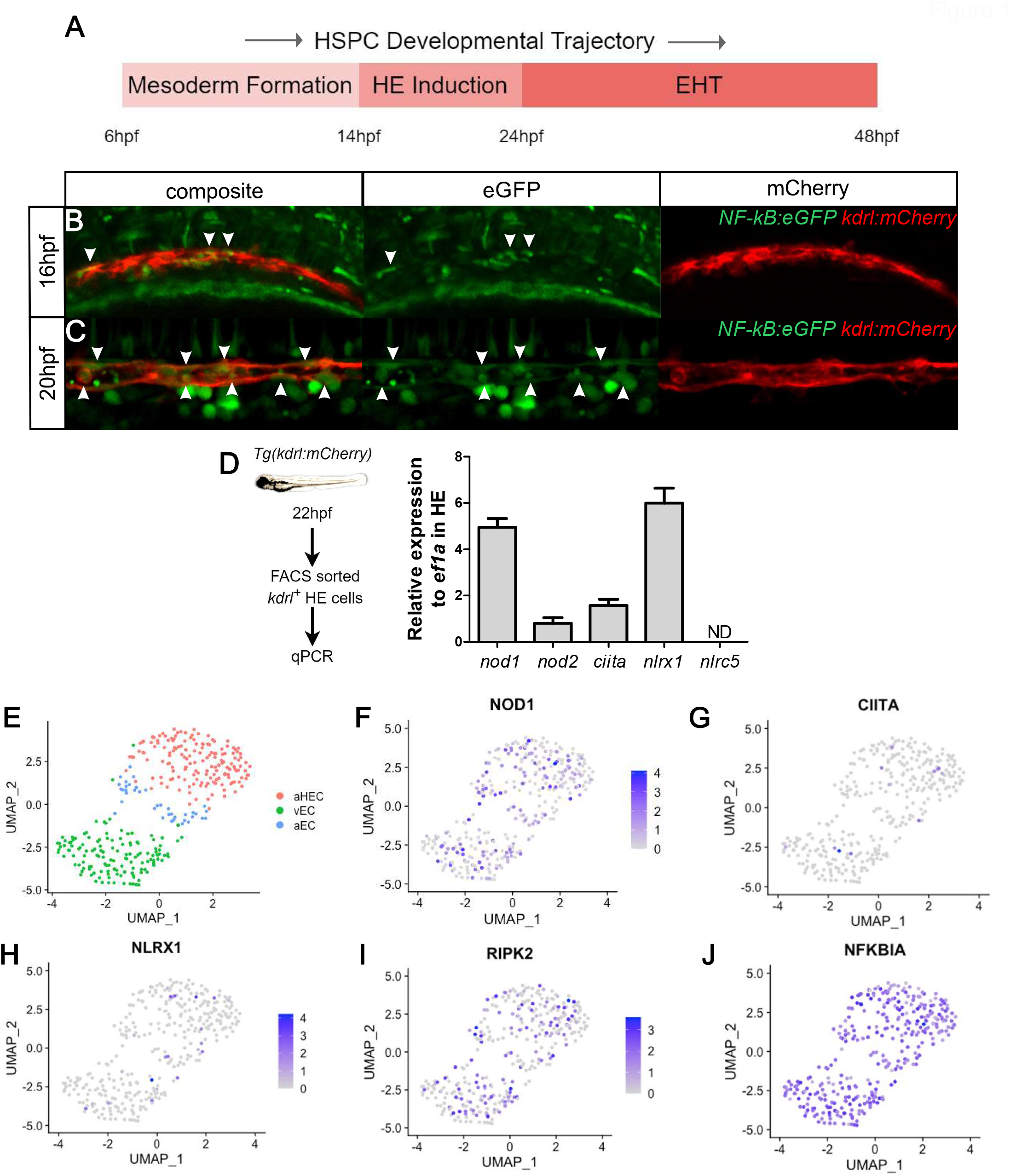

### Nod1 is essential during HSPC specification

Since *NOD1* is one of the most expressed NLRs by the HE in zebrafish and human embryos (Fig. 1D-E), and it can regulate adult HSPC function during bacterial infections (Burberry et al., 2014), as well as induce inflammation in a NF-kB dependent manner, we hypothesized that NOD1 might contribute to HSPC development through the induction of NF-kB in the HE. To test this hypothesis *in vivo*, we performed loss-of function (lof) experiments for Nod1 and directly visualized emerging HSPCs from the floor of the DA in *kdrl:mcherry; cd41:eGFP* double transgenic embryos at 48hpf by confocal microscopy (Fig. 2A-F). The number of double positive *kdrl*^*+*^; *cd41*^*+*^ HSPCs in the floor of the DA was significantly reduced in a dose dependent manner when compared to control embryos when using the Nod1 specific inhibitor Nodinitib-1 (Correa et al., 2011) (Fig. 2A-B), two specific Nod1 antisense morpholinos (MOs), denoted Nod1MO1 and Nod1MO2 (Fig. 2C-D, Fig. S1A-D), and a *nod1*-directed gRNA (Fig. 2E-F, Fig. S1E). In the zebrafish embryo, HSPCs can be visualized along the vDA by expression of *cmyb* using whole-mount *in situ* hybridization (WISH) (Burns et al., 2005). Moreover, the number of *runx1*^*+*^ developing HSPCs in or near the vDA was significantly reduced in *nod1*^*-/-*^ zebrafish null mutants compared to *nod1*^*+/+*^ *siblings*^*-*^ (Fig. 2G-H), validating our previous results. Importantly, 28 hpf *nod1*^*-/-*^ embryos lacked defects on vascular integrity and arterial specification as assessed by WISH for *kdrl* and *efnb2a* (Fig. S1F). Overall, these results robustly show a specific role for Nod1 signaling during HSPC specification.

**Figure 2.**
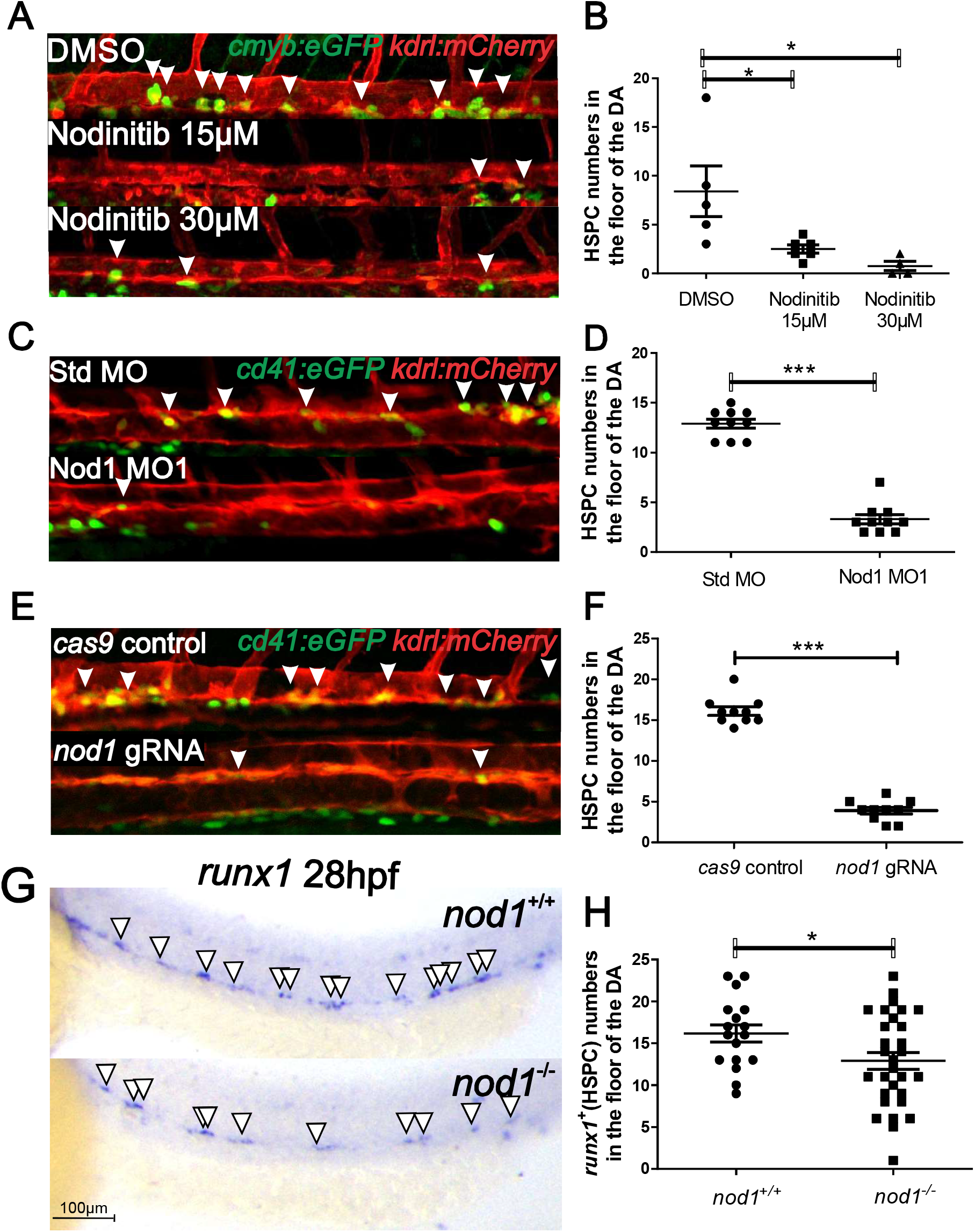

### Nod1 is an early hemogenic endothelial fate inductor

Since *nod1*^*-/-*^ embryos had normal vasculature and arterial development compared to their *nod1*^*+/+*^ counterparts (Fig. S1F), we hypothesized that Nod1 was essential during either HE induction or EHT. In the zebrafish, HE induction occurs between 14-24 hpf, and EHT during 24-48hpf (Bertrand et al., 2010) (Fig. 3A). The number of *cmyb*^*+*^; *kdrl*^*+*^ HSPCs in embryos treated with the Nod1 inhibitor Nodinitib-1 from 24-48 hpf was similar to vehicle-treated animals (Fig. 3A-C). In contrast, HSPC numbers diminished 5-fold if Nod1 was inhibited during HE induction (16-24 hpf) (Fig. 3A-C). These results indicate that Nod1 acts as an early inductor of the HE fate. To validate our conclusion, we next challenged the embryos with C12-iE-DAP, an acylated derivative of the Nod1 agonist iE-DAP that stimulates Nod1 100-1000 times more efficiently (Chamaillard et al., 2003; Park et al., 2007), and performed WISH for *runx1* at 30 hpf. As expected, hyperactivation of Nod1 with its ligand significantly increased the number of early *runx1*^*+*^ HSPCs in the aortic floor (Fig. 3D-E). To confirm the phenotypic specificity of C12-iE-DAP treated embryos, and that the early HSPCs specified in its presence did not lose their identity, we performed WISH for the late HSPC marker *cmyb*, and found a significant and dose-dependent effect upon Nod1 activation (Fig. 3F-G). Together, these results indicate that Nod1 is required to induce the HE during HSPC development.

**Figure 3.**
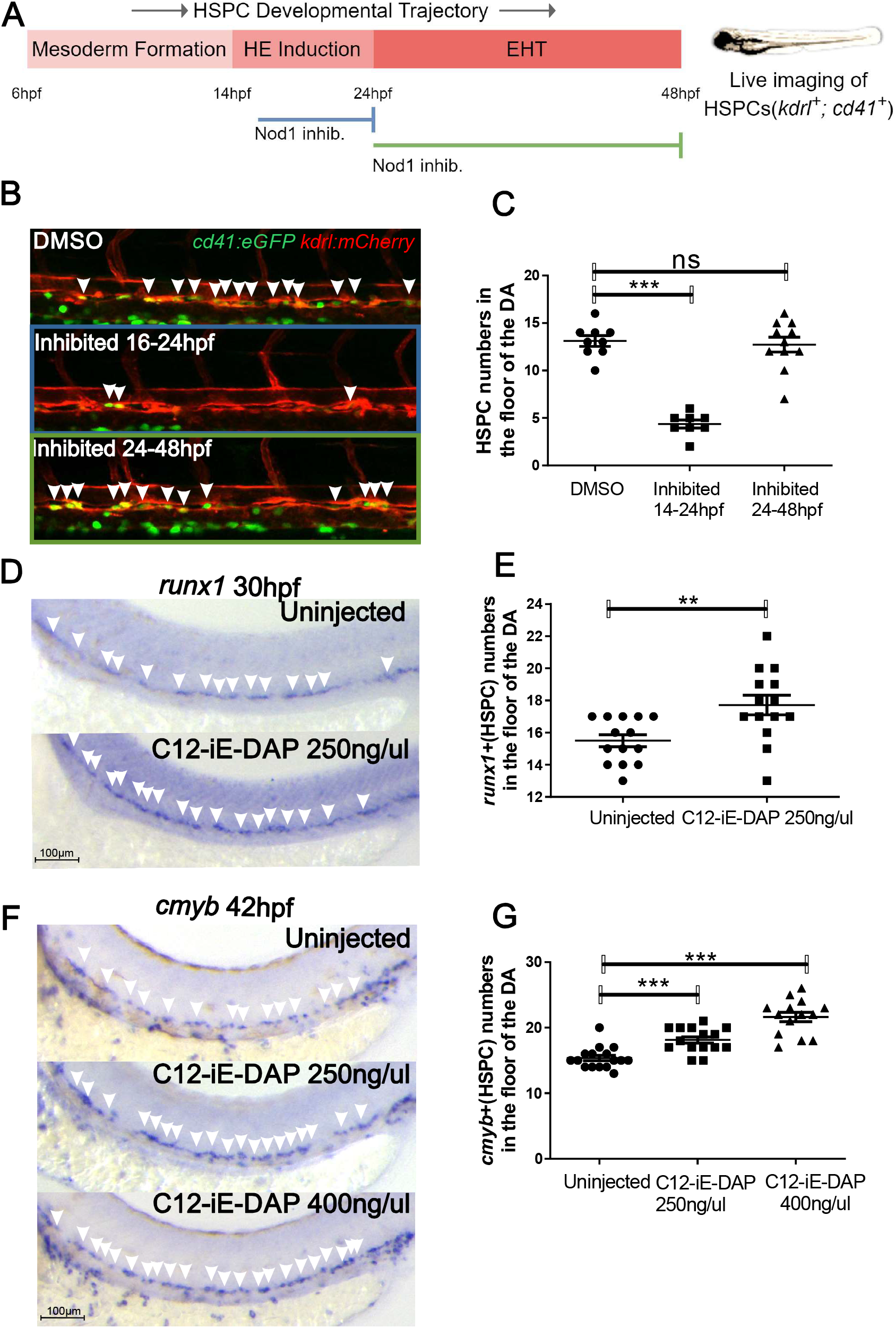

### The effect of Nod1 on HSPC development is Ripk2-dependent

In classical inflammation, Nod1 resides in an autoinhibited monomeric state in the cytosol. Upon peptidoglycan sensing, Nod1 transitions to an open conformation, which allows self-oligomerization and recruitment of its main effector kinase receptor-interacting serine/threonine-protein kinase 2 (Ripk2) through homotypic CARD-CARD interactions (Caruso et al., 2014). Since the canonical adaptor protein required for the activation of the Nod1 signaling pathway is Ripk2 during infection, we queried if signaling through Nod1 similarly activated Ripk2 to specify HSPCs in the aseptic embryo. First, we quantified live HSPC numbers by confocal imaging of *Tg(kdrl:mcherry*^*+*^, *cd41:eGFP*^*+*^) embryos after Ripk2 depletion by a specific translating blocking morpholino, and found a significant decrease compared to control embryos (Fig. 4A-B). To validate this result, we performed WISH for *runx1* (30 hpf) and *cmyb* (42 hpf) in published *ripk2* zebrafish mutants (Jurynec et al., 2018). As expected, both *runx1*^*+*^ and *cmyb*^*+*^ cells were significantly downregulated in *ripk2*^*-/-*^ embryos compared to wildtype (wt) controls (Fig. 4C-D), faithfully mimicking the lack of developing HSPCs observed upon Nod1 depletion. If Ripk2 activation was indeed required downstream of Nod1 function for HSPC specification, then ectopic expression of the hyperactive *ripk2*^*104Asp*^ (Jurynec et al., 2018) by mRNA injection into Nod1-deficient zygotes should rescue the lack of HSPCs. Accordingly, enforced *ripk2*^*104Asp*^ expression rescued *runx1*^*+*^ HSPC numbers in Nod1 morphants, indicating that Nod1 signals through Ripk2 to specify the HE (Fig. 4E-F). To ensure that loss of HSPCs in the absence of Ripk2 was not due to impaired vascular and arterial development, we performed WISH for the endothelial and arterial markers *kdrl* and *efnb2a* (Lawson et al., 2001), respectively. No vascular or arterial abnormalities were observed in *ripk2*^*-/-*^ compared to wt control embryos (Fig. S2A). Nod1 and Ripk2 both contain CARD domains, which are able to directly interact with caspases (Park, 2019). While there is no direct link between Nod signaling and apoptosis (Heim et al., 2019), it has been shown that NOD1 can activate Caspase-9 in a RIPK2-dependent manner (Inohara et al., 1999). To address if the loss of HSPCs in *ripk2*^*-/-*^ embryos was due to the apoptosis of endothelial cells, we performed a Terminal deoxynucleotidyl transferase dUTP nick end labeling (TUNEL) assay. Analysis of the aorta-gonad-pronephros (AGP) region by confocal microscopy in *ripk2*^*-/-*^ embryos at 18 hpf and 23 hpf showed no increased apoptosis within the aorta-gonad-pronephros (AGP) (Fig. S2B-E), the fish analog of the mammalian aorta-gonad-mesonephros. Finally, we examined T cell development using *lck:eGFP* transgenic animals (Langenau et al., 2004) at 5 dpf in the absence of Nod1 or Ripk2, and found lack of T cells (Fig. 4G), supporting a role for Nod1-Ripk2 signaling in the specification of definitive hematopoietic progenitors. Together, these findings indicate that decreased HSPC numbers in Nod1- and Ripk2-deficient embryos is not caused by apoptosis, but due to failure in early patterning of definitive HE.

**Figure 4.**
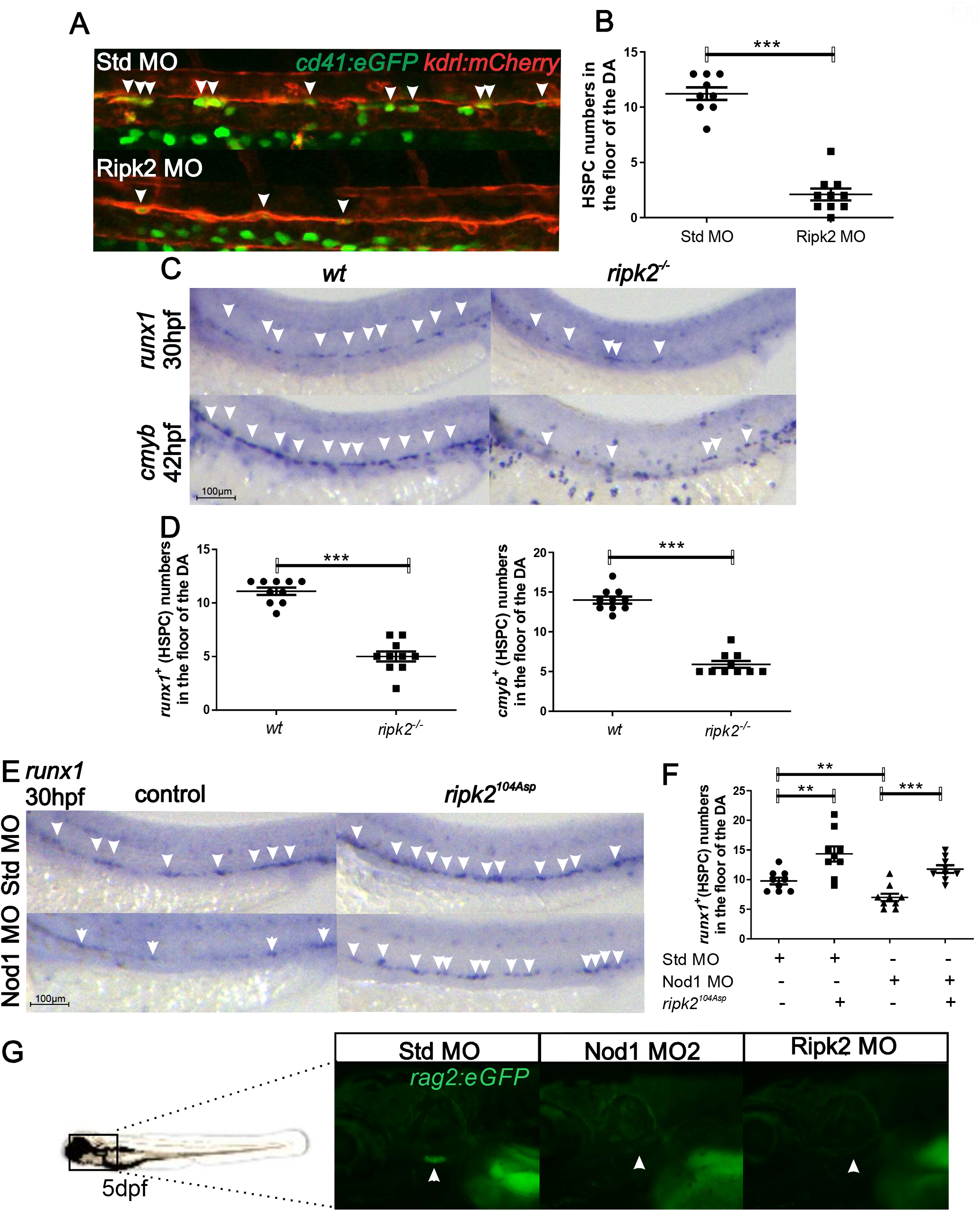

### Transcriptomic analysis of Nod-deficient HE identifies deregulated hematopoietic and immune programs

To identify molecular effectors that were defective within the HE in the absence of Nod1, we purified by FACS *kdrl*^*+*^ HE at 22 hpf from Nod1 morphants and Std-MO injected controls and performed RNA-sequencing (RNA-seq) (Fig. 5A). As expected, a sample correlation heatmap showed the strongest correlation among the Nod1-MO and control Std-MO triplicates Fig. S3A. The MA plot showed the number of differentially expressed genes (DEGs) between Nod1-deficient HE compared to Std control (Fig. 5B), with 797 down-regulated and 792 up-regulated genes between both conditions with an adjusted p-value *P*_adj_ value ≤ 0.05 (Fig. 5B-C). The top 100 DEGs are represented in the heatmap in Fig. S3B. GO enrichment analysis for biological processes in down-regulated DEGs included “Immune System Process”, “Erythrocyte Differentiation”, “Myeloid Cell Differentiation” and “Embryonic Hemopoiesis” (Fig. 5E-H). In contrast, MAPK pathways were unaffected (not shown), suggesting the failure to establish the hematopoietic program in the absence of Nod1 was through an immune-related mechanism rather than the MAPK pathway.

**Figure 5.**
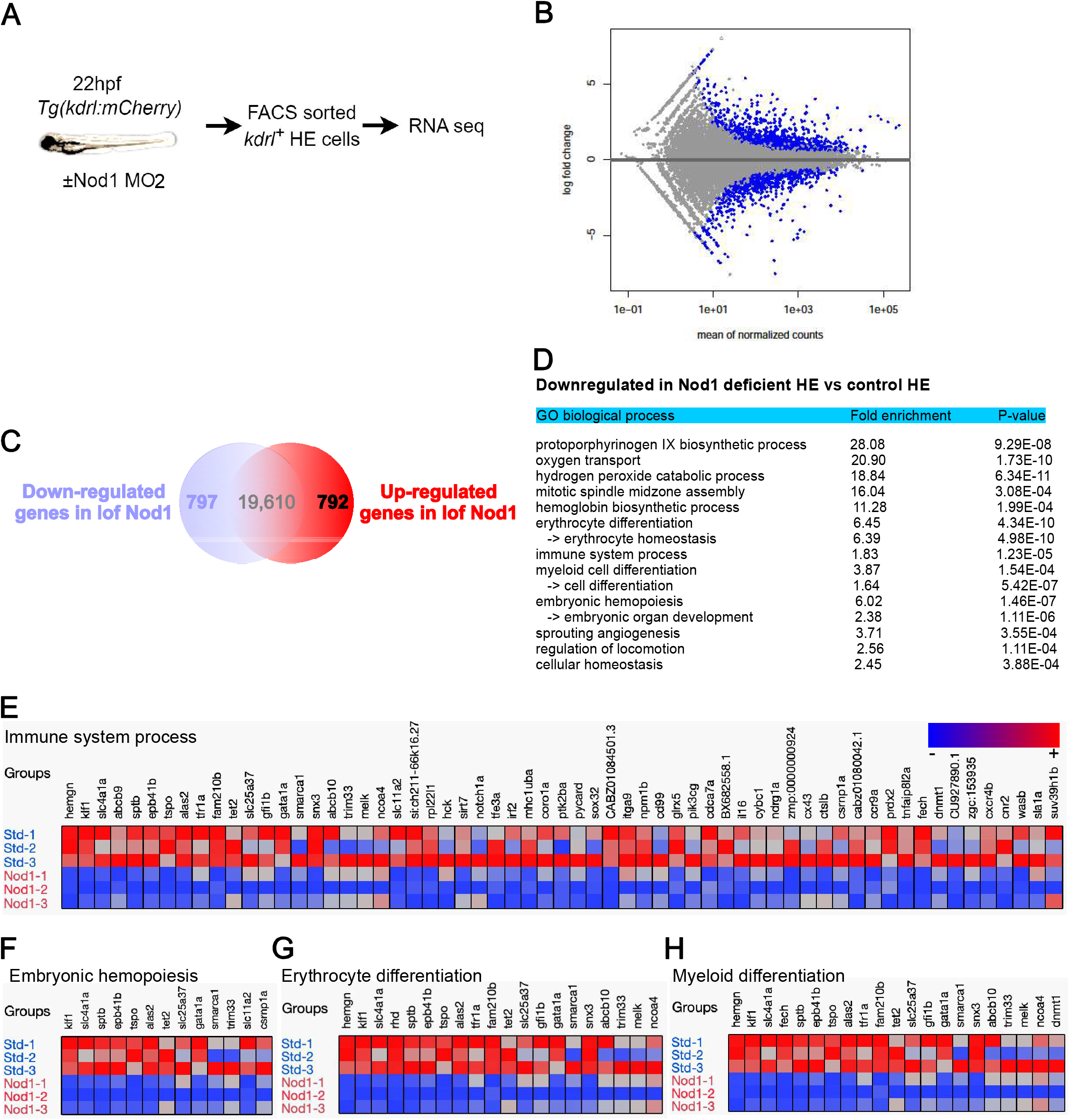

### Nod1 and Ripk2 signal through NF-kB to specify HSPCs

NF-kB is the master transcription factor of inflammation (Capece et al., 2022; Espin-Palazon and Traver, 2016), and its activation is required to specify HSPCs in the aseptic vertebrate embryo (Espin-Palazon et al., 2014; He et al., 2015). Moreover, during bacterial infections NOD1 and RIPK2 activate NF-kB after sensing peptidoglycan. These lines of evidence suggested that NOD1 might activate NF-kB during HE induction to specify HSPCs. To address this, we performed live confocal imaging of *Tg(kdrl:mcherry; NF-kB:eGFP)* in Nod1 lof experiments. As anticipated, we observed a remarkable down-regulation of NF-kB activity in the HE at 22 hpf in Nod1-deficient embryos compared to control siblings (Fig. 6A). The number and levels of NF-kB activity within the HE (*kdrl*_*+*_) were quantified by flow cytometry, showing a significant reduction in both (Fig. 6B). If NF-kB signaling was indeed required downstream of Nod1 and Ripk2 function for HSPC specification, then ectopic expression of a constitutively activated Inhibitor of Nuclear Factor Kappa B Kinase Subunit Beta (Ikkb), the main kinase activating canonical NF-kB that phosphorylates the inhibitor in the inhibitor/NF-kB complex, causing activation of NF-kB (Espin-Palazon and Traver, 2016) (Fig. S4), should rescue the lack of HSPCs in nod1- or ripk2-deficient embryos. In order to activate NF-kB, IKKB undergoes phosphorylation on residues Ser-177/181 located in the amino-terminal activation T loop (Fig. S4). Substitution of these two serine residues to negatively charged glutamic acids renders human IKKB constitutively active (Sasaki et al., 2006). We hypothesized that substitution of the highly conserved Ser-177/181 by Glu in the zebrafish Ikkb (herein called constitutively activated ikkb, Ikkb_CA_) would also result in its hyperactivation. mRNA injection of Ikkb_CA_ into one-cell stage *NFkB:eGFP* embryos increased NF-kB activation as compared to Ikkb control injection (Figure 6C), validating this approach. We then performed rescue experiments by mRNA overexpression of Ikkb_CA_, or Ikkb control, in Ripk2-deficient embryos and performed WISH for *cmyb* at 42 hpf. As shown in Fig. 6D-E, enforced activation of NF-kB completely rescued HSPC numbers in Ripk2 morphants, demonstrating that Nod1/Ripk2 signaling activates the NF-kB pathway within HE to drive HSPC fate.

**Figure 6.**
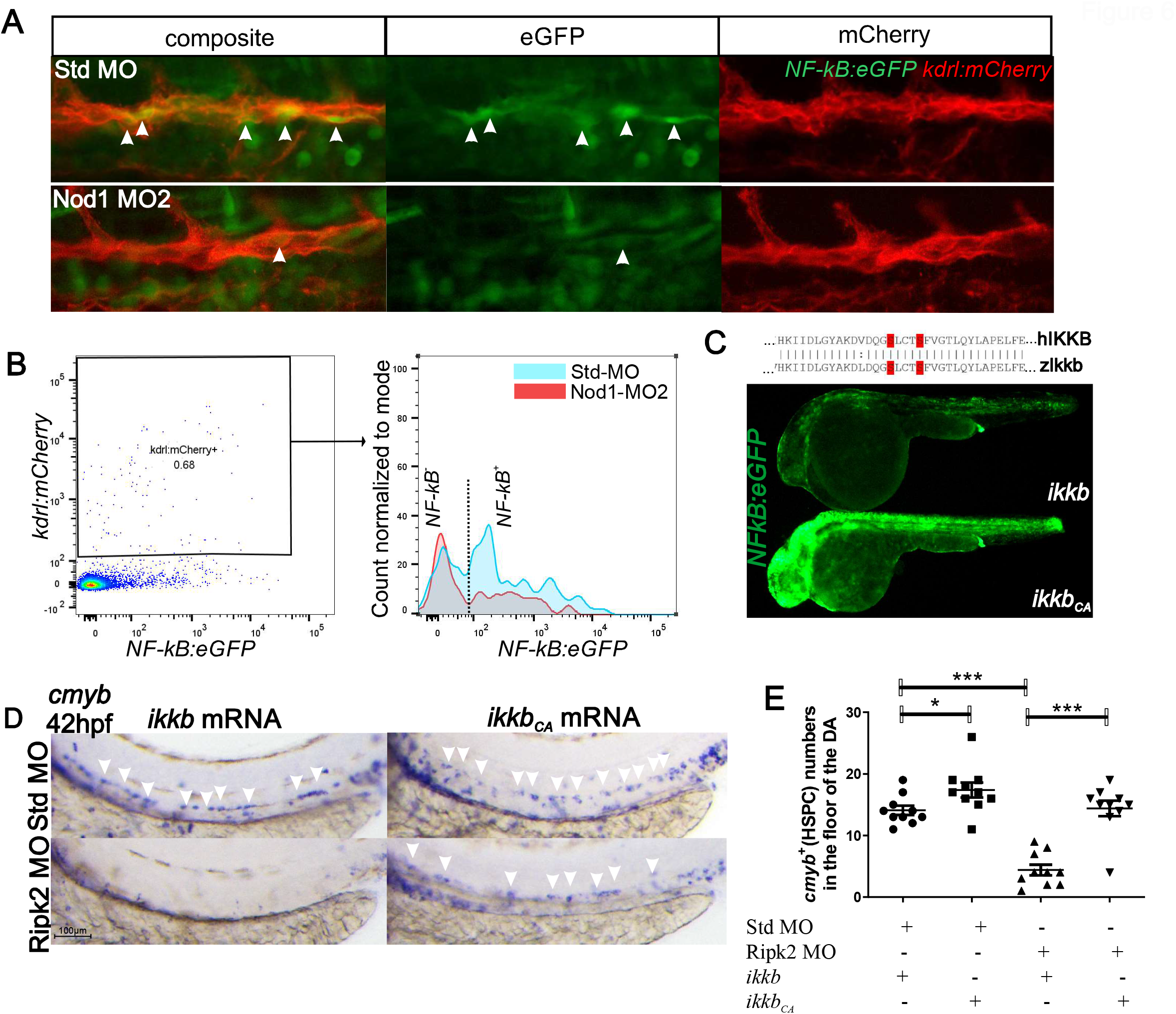

### Rho-GTPases power the Nod1-Ripk2-NF-kB axis in vivo to specify HSPCs

It was unexpected that the Nod1-Ripk2-NF-kB axis, a key pathway underlying the canonical sensing of PAMPs, was required to generate the founders of the adult hematopoietic system during natural embryonic development. Since the emergence of HSPCs from the aortic floor occurs in the aseptic embryo, whether it is *in utero* in mammals or within the chorion in teleosts, we hypothesized that other stimuli beyond PAMPs could be activating Nod1 in the context of developmental inflammation. Because it has been shown that NOD1 senses cytosolic microbial products by monitoring the activation state of small Rho GTPases (Keestra et al., 2013), and Rho GTPases can also regulate hematopoietic stem cell functions (Cancelas and Williams, 2009) and promote HSPC formation *in vivo* (Lundin et al., 2020), we postulated that Rho-GTPases activate Nod1 during HSPC specification. We first queried if *rac1, cdc42*, and *rho*, the three small Rho GTPases that can activate Nod1 in response to pathogens (Keestra et al., 2013), were expressed during HE induction. qPCR from FACS isolated *kdrl:mcherry*^+^ HE showed that *rac1a/b* and *cdc42* were expressed by the HE, but not *rho* (Fig. 7A). Treatment of *Tg(kdrl:mcherry; Cd41:eGFP)* embryos with the small G-protein inhibitors hydrochloride, Rho kinase inhibitor III, and ML141, which specifically inhibit Rac1, Rho kinase, and CDC42 (Table S3), respectively, from 16 hpf reduced 10-fold the number of *kdrl*^*+*^, *cd41*^*+*^ HSPCs by live confocal imaging at 48 hpf (Fig. 7B-C). Furthermore, enforced NF-kB activation by *ikkb*_*CA*_ overexpression significantly rescued the phenotypic levels of *kdrl*^*+*^, *cd41*^*+*^ HSPCs in the vDA upon Rac1 inhibition, and partially upon Cdc42 inhibition, but not after Rho kinase inhibition (Fig. 7C). These findings suggest that Rac1 specifies HSPCs through the activation of pro-inflammatory signaling. To confirm that Rac1 is essential during HSPC specification, we next injected a previously validated Rac1-specific morpholino (Table S2) that targets both zebrafish ohnologues (Rac1a and Rac1b), and quantified HSPCs by confocal microscopy. As shown in Fig. S5, zebrafish Rac1a and Rac1b are highly conserved with human and mice RAC1, with 99% and 98% identity, respectively (Fig. S5A). *Kdrl*^*+*^, *cd41*^*+*^ HSPC numbers were significantly reduced after Rac1a/b depletion. Overall, these results indicated that the small Rho GTPase Rac1 activates Nod1-Ripk2-NF-kB signaling intrinsically within the HE to specify HSPCs.

**Figure 7.**
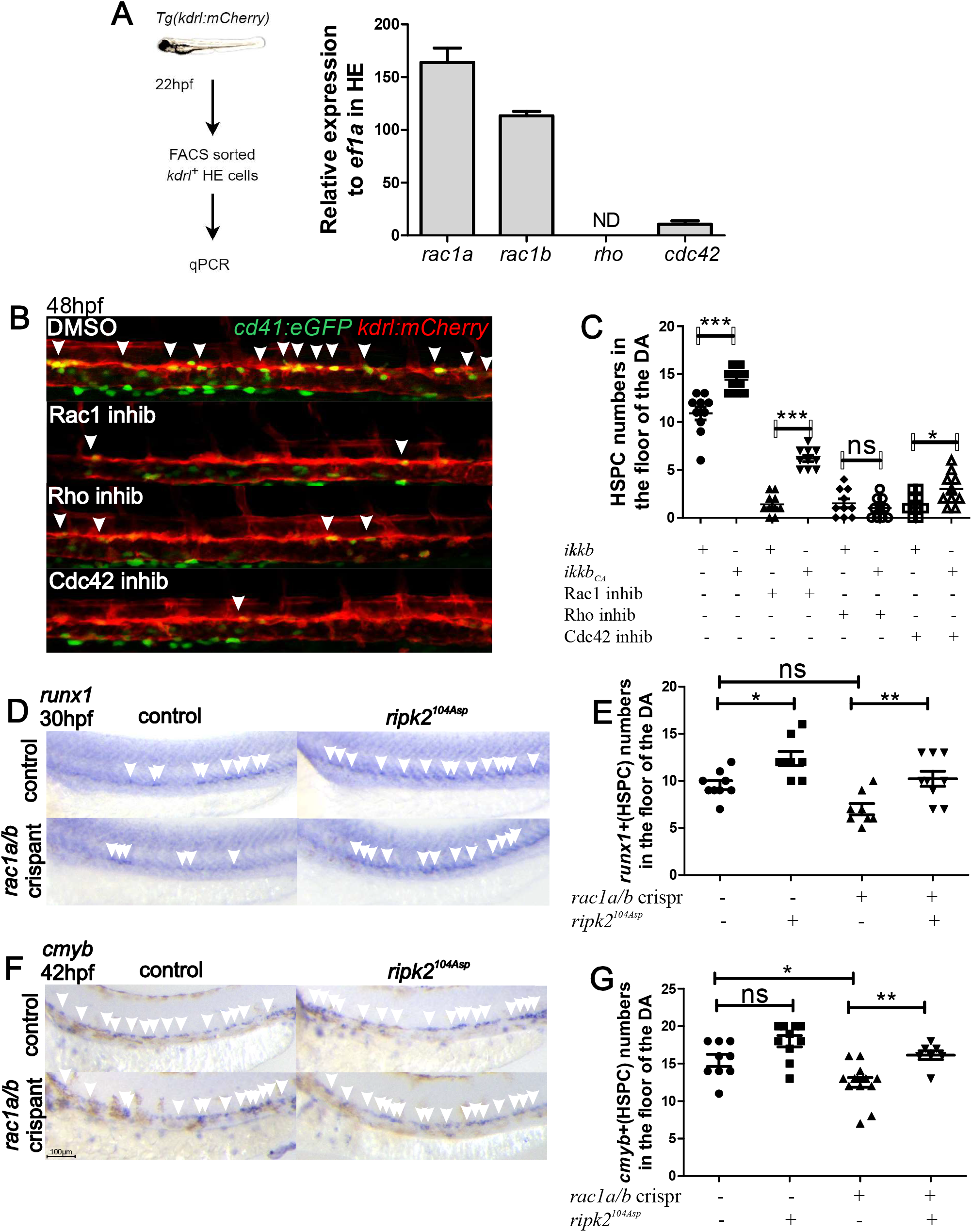

### The function of Nod1 is conserved in the development of definitive human HSPCs

Our final goal was to address whether NOD1 plays similar roles in human development hematopoiesis. For this purpose, first we queried if the human versions of the identified genetic axis Rac1-Nod1-Ripk2 were expressed in iPSC-derived human HSPCs. Transcriptomic data at the single cell resolution from definitive uncommitted hematopoietic progenitors and primed differentiated hematopoietic lineages (Fidanza et al., 2020) showed that as expected, human *NOD1* transcripts were enriched in the endothelial and HSC/MPP clusters (Fig. 8A). In addition, *RIPK2, RAC1, and NFKBIA* were expressed throughout all clusters (Fig. 8A). Next, we assessed the effect of NOD1 deficiency on the hematopoietic potential of embryonic bodies treated with CHIR to induce definitive hematopoiesis (Fidanza et al., 2020; Sturgeon et al., 2014). We incubated the cells either from day 2, or day 8, with the NOD1 inhibitor Nodinitib-1 (iNOD1), and quantified the number of hematopoietic progenitors at day 15 after plating 20K CD34^+^ cells on OP9 co-culture from day 8 (Fig. 8B). The number of hematopoietic progenitors in suspension dropped 6-fold compared to vehicle-treated control when NOD1 was inhibited from day 2. In contrast, no significant differences were found when NOD1 function was inhibited during EHT (Fig. 8B-D). Cell viability was similar between control and iNOD1-treated cells from day 2 (Fig. S6A-B), but the percentage of CD34_high_ endothelial cells was significantly higher (Fig. S6C-D), indicating that the defect on hematopoietic specification in the absence of NOD1 was specific, and loss of NOD1 favors a non-HE endothelial fate. Altogether, these data show that NOD1 signaling is a conserved mechanism during vertebrate development that patterns the HE for their subsequent trans-differentiation into definitive hematopoietic progenitors (Fig. 8E).

**Figure 8.**
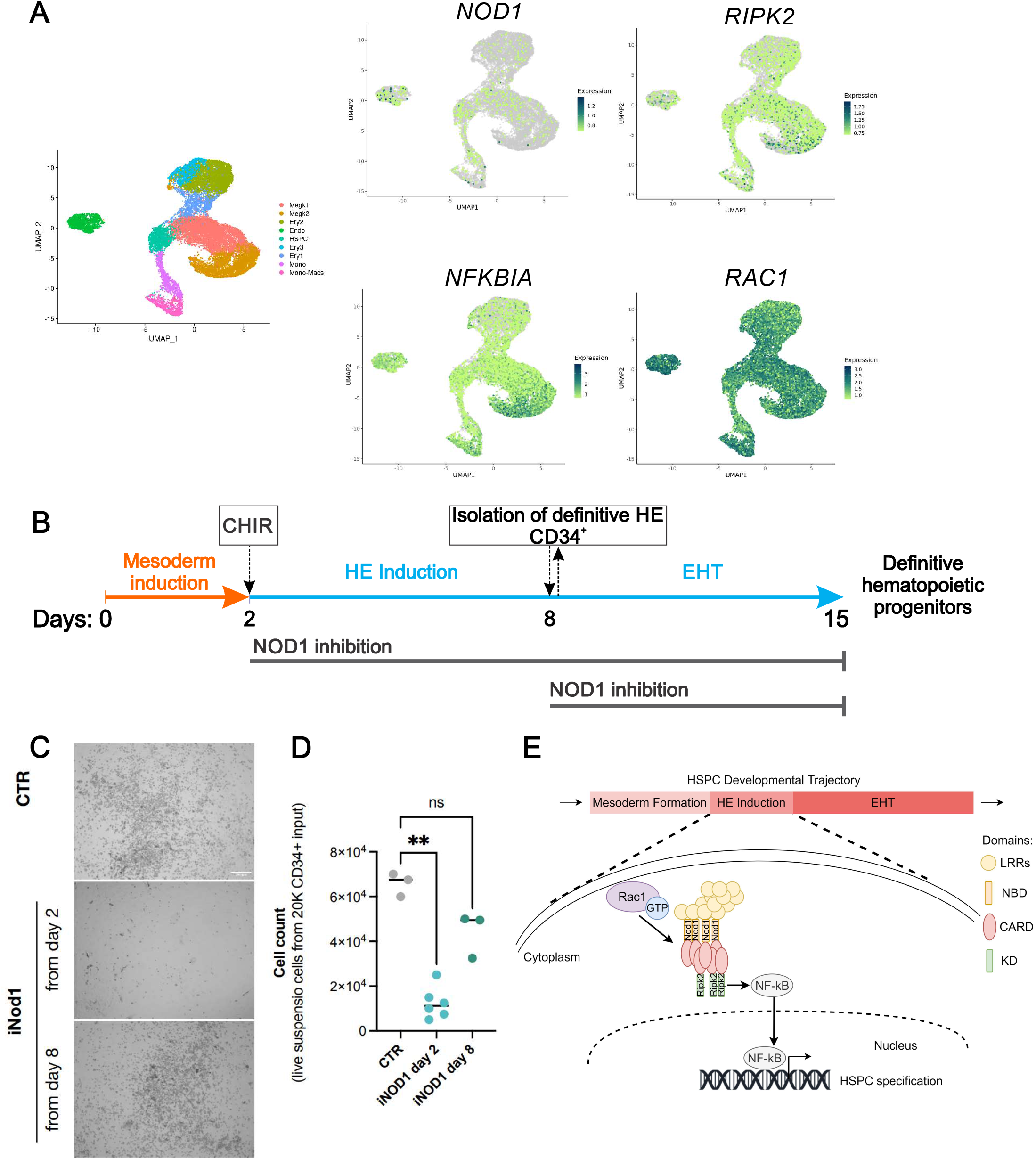

## Discussion

The capability to specify HSPCs from hPSCs will require an in-depth understanding of the natural molecular mechanisms utilized by the developing embryo to generate HSPCs. Here, we identified *in vivo* a previously overlooked Nod1-dependent mechanism that intrinsically initiates inflammatory signaling in the HE, resulting in its proper induction and subsequent HSPC specification. This signaling pathway is the earliest identified activator of inflammation required for HSPC fate. First, we identified that non-inflammasome forming NLRs, including *NOD1* were expressed during HE induction in zebrafish and human embryos. Second, we demonstrated that Nod1 and Ripk2 worked in a linear genetic pathway through NF-kB activation to establish HE competency. Third, we showed that this mechanism is highly conserved during human definitive hematopoietic development. Lastly, our data revealed that small Rho GTPases, especially Rac1, were the source of Nod1 activation required to pattern the HE. Together, these findings support a highly conserved model of vertebrate hematopoietic development in which small Rho GTPases and pattern recognition receptors orchestrate the activation of the master inflammatory transcription factor NF-kB in the HE to facilitate definitive hematopoiesis (Fig. 8E).

The idea that a classical signaling pathway underlying the canonical sensing of pathogenic peptidoglycan is required in the aseptic embryo to generate the founders of the adult hematopoietic system was unexpected, since the emergence of HSPCs from the aortic floor occurs in the absence of pathogens. Our study identifies for the first time that the non-inflammasome forming NLRs are fundamental inflammatory mediators that initiate the earliest NF-kB induction in the HE to drive HSPC fate. Hyperactivation of Nod1, Ripk2, or NF-kB in this work led to a consistent increased on early specified HSPC numbers, indicating that this inflammatory signaling pathway acts inducing HE rather than its maintenance or survival.

Although recent work demonstrated that NOD1 can also sense cellular perturbations like changes on actin cytoskeleton (Bhavsar et al., 2013; Boyer et al., 2011; da Silva Correia et al., 2007; Keestra et al., 2013; Kufer et al., 2008; Mayor et al., 2007; Uehara et al., 2006) or endoplasmic reticulum (ER) stress (Bernales et al., 2006; Celli and Tsolis, 2015; Keestra-Gounder et al., 2016; Keestra-Gounder and Tsolis, 2017; Ogata et al., 2006), these perturbations were caused by pathogen disruptions. Here, we provide the first evidence that in the context of embryonic development, small Rho GTPases can induce Nod1 signaling in a pathogen-independent manner. Whether changes in the cytoskeleton or ER stress occur naturally in the endothelial cells that transition to HE fate, and these stresses are captured by small Rho GTPases to activate Nod1, would need further assessment. Our data showing that Rac1 is essential during early HSPC genesis supports previous studies in mice where depletion of Rac1 resulted in the absence of intra-aortic clusters (Ghiaur et al., 2008). However, our study provides the molecular mechanism by which Rac1 drives HSPC specification, connecting for the first time the function of Rho GTPases through the Nod1-Ripk2 inflammatory route. Our expression and rescue data showing high correlation among the type of Rho GTPase expressed in the HE and efficacy of the rescue indicated that Nod1 could potentially be activated by several small Rho GTPases, but the specificity could be due to the expression levels of these small Rho GTPases in the HE. In addition, previous studies in mice and zebrafish also demonstrated that Rho GTPases are essential for HSPC specification and migration after the initiation of blood flow (Ghiaur et al., 2008; Lundin et al., 2020). Lundin et al. demonstrated that mechanical forces induced by blood flow stimulated Rho-GTPase function to activate Yap-mediated HSPC specification *in vivo*. Therefore, it is clear that Rho-GTPases are key modulators of HSCP specification in a multistep fashion. Studying the perturbations that HE is subjected to during its induction could provide important insights on the mechanisms that trigger Rac1-Nod1-Ripk2-NFkB during HE induction.

Although it has been shown that primitive myeloid cells are an important source of pro-inflammatory cytokines that drive developmental inflammation and HSPC fate (Espin-Palazon et al., 2014; Sawamiphak et al., 2014; Tie et al., 2019), our work here uncovered a much earlier inflammatory requirement that primes HE patterning for definitive hematopoiesis, suggesting a multi-step requirement for inflammatory signaling during the ontogeny of HSPCs. Since the HE niche is deprived of primitive myeloid cells before the onset of blood flow, it is plausible that different triggers might contribute to pro-inflammatory signaling during HSPCs specification, being the Nod1 pathway one of the first endogenous initiators. It is noteworthy that Nod1 mutations have been associated with autoimmune disorders (Hysi et al., 2005). Our results demonstrating impaired developmental hematopoiesis in the absence of Nod1 provide the foundation to investigate whether the autoimmune disorders linked to NOD1 mutations could be caused, at least in part, by the defective establishment of the hematopoietic system during embryonic development.

Importantly, the precise temporal identification of the Nod1 requirement *in vivo* was successfully translated *in vitro* in a human system of hematopoietic development, demonstrating once more than animal models such as zebrafish are key to discover novel mechanisms of HSPC fate induction. Since hyperactivation of Nod1, Ripk2, or NF-kB during HE induction in our *in vivo* zebrafish model robustly increased HSPC numbers, the addition of compounds that activate these molecular players during the HE-like induction in *in vitro* protocols of human hematopoietic differentiation could help derive a more competent HE to achieve HSPC fate.

In summary, we have reported a previously overlooked requirement for inflammatory signaling in the early HE induction that regulates HSPC fate, their molecular components, and its temporal specificity. Inducing this molecular mechanism *in vitro* could enhance the derivation of definitive human hematopoietic progenitors and position us closer to achieve the generation of patient-specific HSCs for their use in regenerative medicine.

## Acknowledgments

This work was supported by the NIH-NIDDK R01 (R01DK131162), R03 (R03DK125661), K01 award (7K01DK115661) to R.E-P.; ASH Global Award and EHA Advanced Research Grant to A.F., and the Roy J. Carver Charitable Trust (#21–5532). This publication includes data generated at the UCSD IGM Genomics Center utilizing an Illumina NovaSeq 6000 that was purchased with funding from a National Institutes of Health SIG grant (#S10 OD026929). The authors are indebted to Jeffrey Essner and Maura McGrail for their support with fish husbandry, and Michael Jurynec for kindly donating *ripk2*^*-/-*^ zebrafish mutants and plasmids containing wt and mutant versions of *ripk2*.

## Author contributions

X.C., R.B., E.S., A.F., C.C., and R.E.-P. designed experiments; X.C., R.B., A.G., E.S., A.F., C.C., and R.E.-P. performed research; X.C., R.B., A.G., E.S., A.F., C.C., and R.E.-P. analyzed data; Y.Z. and K.D. analyzed RNA-seq data; and R.E.-P. and X.C. wrote the paper with minor contributions from remaining authors.

