## Supplemental figures for "Nod1-dependent NF-kB activation initiates hematopoietic stem cell specification in response to small Rho GTPases"

Sup 1

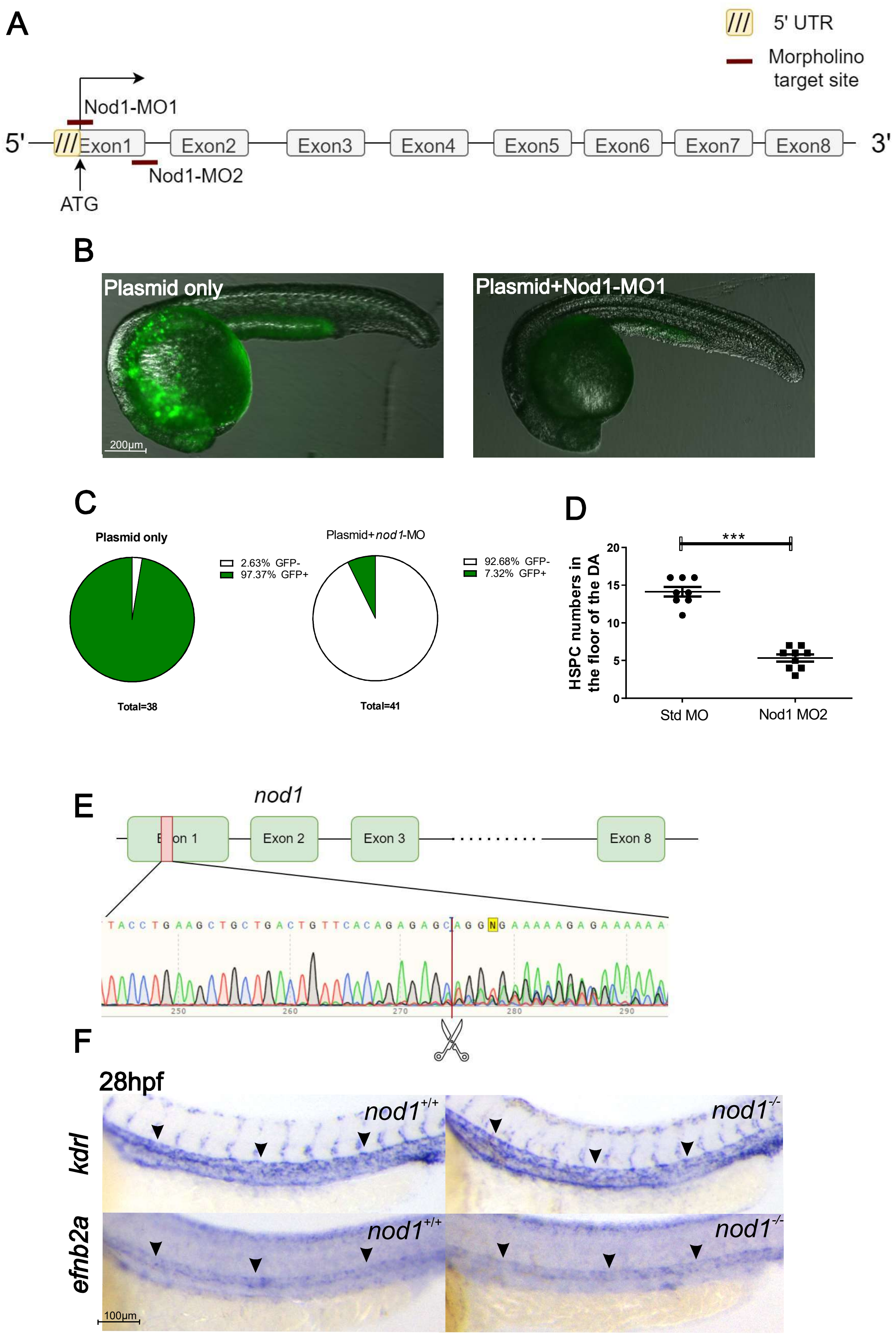

A
5' UTR
Morpholino target site
Nod1-MO1
5'
Exon1
Exon2
Exon3
Exon4
Exon5
Exon6
Exon7
Exon8
3'
Nod1-MO2
ATG
B
Plasmid only
Plasmid+Nod1-MO1
200µm
C
Plasmid only
2.63% GFP-
97.37% GFP+
Total=38
Plasmid+nod1-MO
92.68% GFP-
7.32% GFP+
Total=41
D
HSPC numbers in the floor of the DA
***
20
15
10
5
0
Std MO
Nod1 MO2
E
nod1
Exon 1
Exon 2
Exon 3
Exon 8
TACCTGAAGCTGCTGACTGTTCACAGAGAGCAGGNGAAAAAGAGAAAAAA
250
260
270
280
290
F
28hpf
kdrl
efnb2a
nod1+/+
nod1-/-
100µm

Sup 2

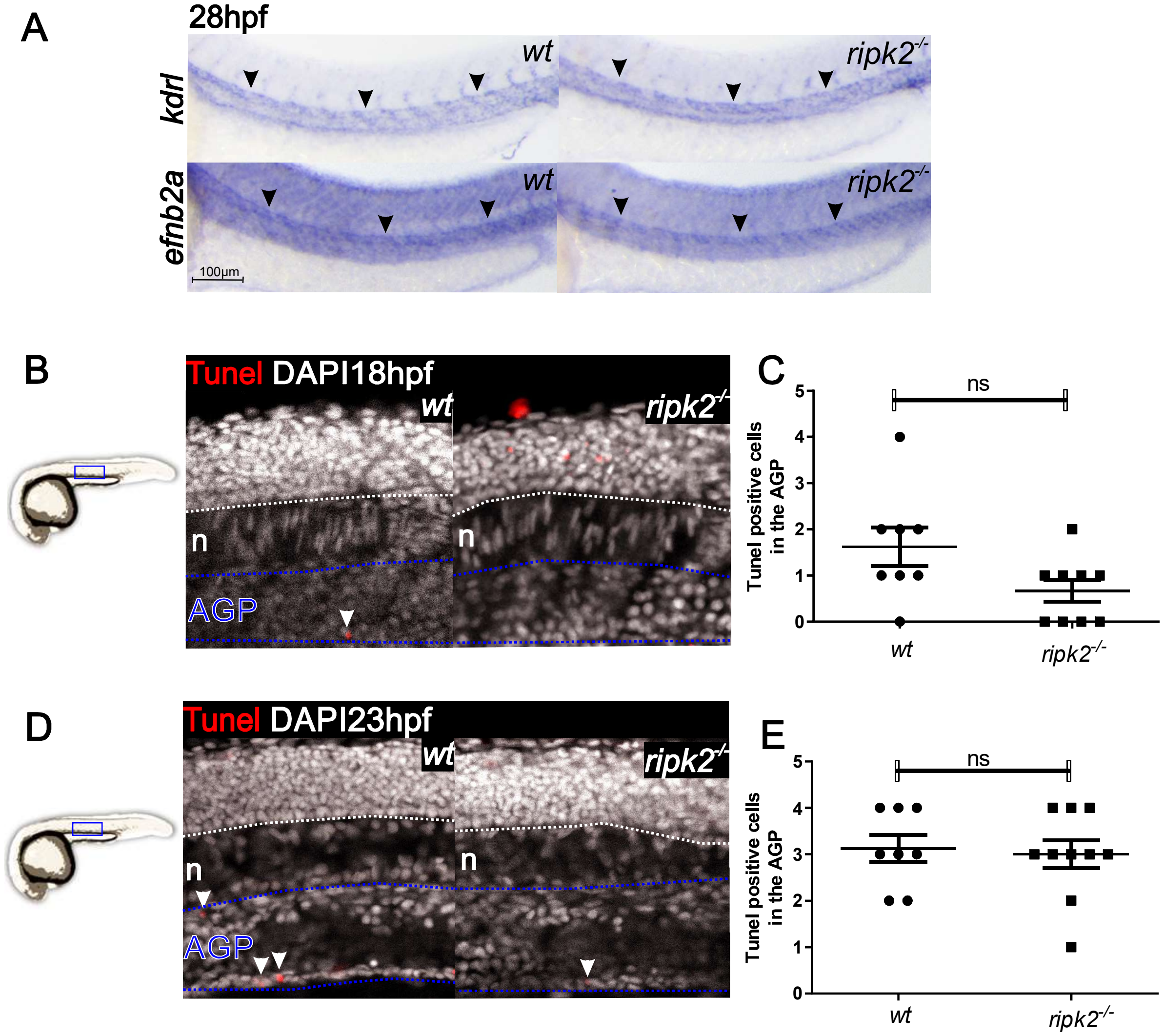
A
28hpf
kdrl
efnb2a
wt
ripk2-/-
100µm
B
Tunel DAPI18hpf
wt
ripk2-/-
n
AGP
C
Tunel positive cells in the AGP
ns
0
1
2
3
4
5
wt
ripk2-/-
D
Tunel DAPI23hpf
E

Sup 3

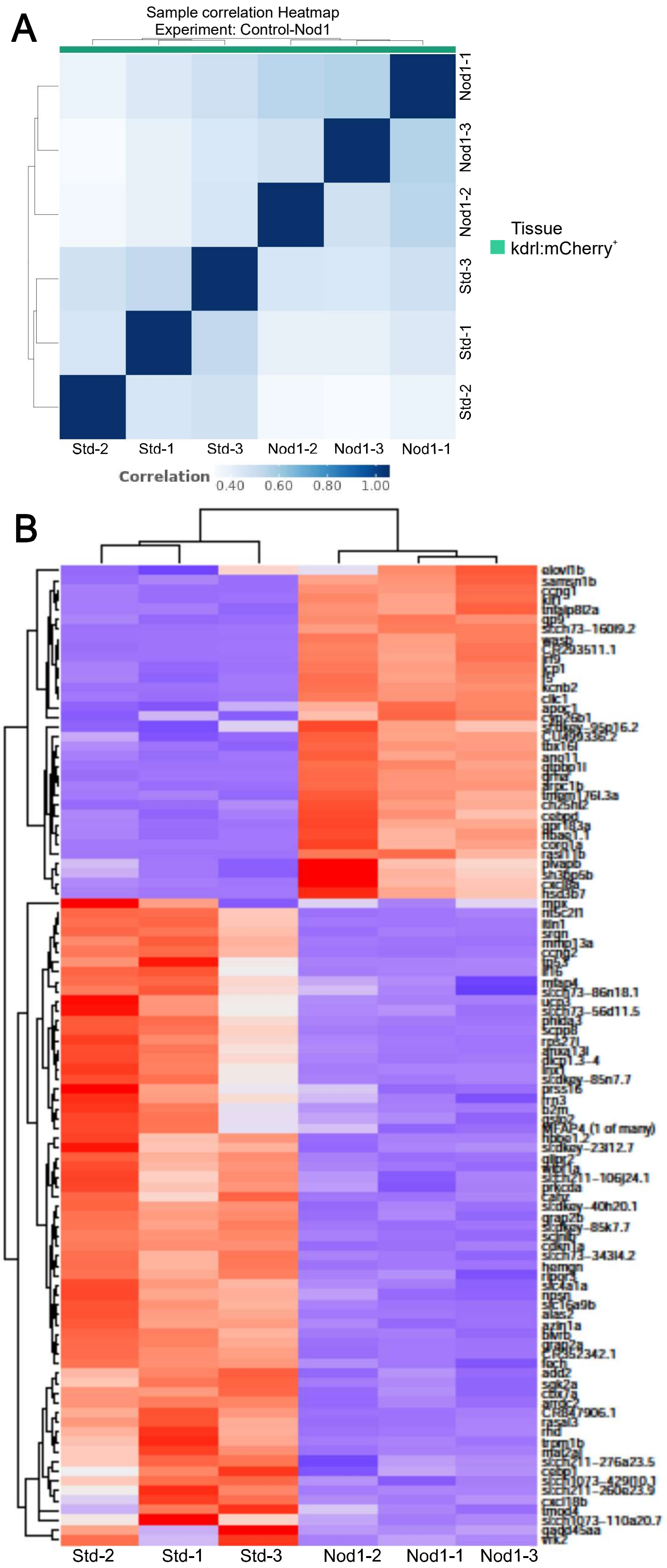
A
Sample correlation Heatmap
Experiment: Control-Nod1
Nod1-1
Nod1-3
Nod1-2
Std-3
Std-1
Std-2
Std-2
Std-1
Std-3
Nod1-2
Nod1-3
Nod1-1
Tissue
kdrl:mCherry+
Correlation
0.40
0.60
0.80
1.00
B
Std-2
Std-1
Std-3
Nod1-2
Nod1-1
Nod1-3

Sup 4

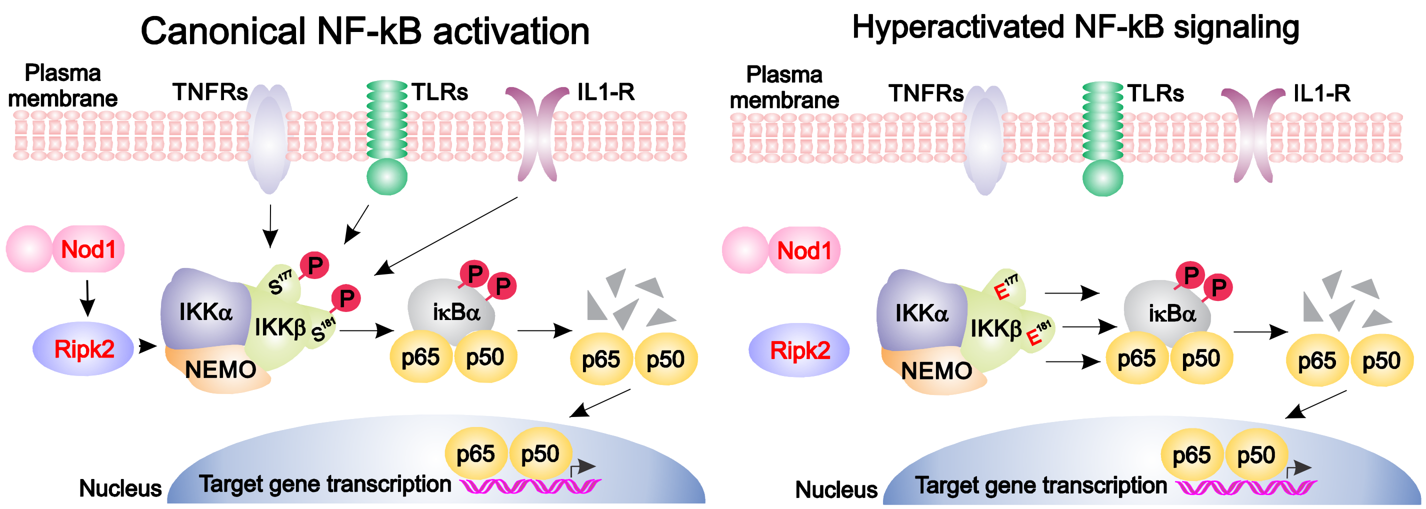
Canonical NF-kB activation
Plasma membrane
TNFRs
TLRs
IL1-R
Nod1
Ripk2
IKKα
IKKβ
NEMO
S177
S181
P
IκBα
p65
p50
Nucleus
Target gene transcription
Hyperactivated NF-kB signaling
E177
E181

Sup 5

A

Rac1a: Protein similarity: 99%

```
hRac1     MQAIKCVVVGDGAVGKTCLLISYTTNAFPGEYIPTVFDNYSANVMVDGKPVNLGLWDTAG   60
mRac1     MQAIKCVVVGDGAVGKTCLLISYTTNAFPGEYIPTVFDNYSANVMVDGKPVNLGLWDTAG   60
zRac1a    MQAIKCVVVGDGAVGKTCLLISYTTNAFPGEYIPTVFDNYSANVMVDGKPVNLGLWDTAG   60
          ************************************************************

hRac1     QEDYDRLRPLSYPQTDVFLICFSLVSPASFENVRAKWYPEVRHHCPNTPIILVGTKLDLR   120
mRac1     QEDYDRLRPLSYPQTDVFLICFSLVSPASFENVRAKWYPEVRHHCPNTPIILVGTKLDLR   120
zRac1a    QEDYDRLRPLSYPQTDVFLICFSLVSPASFENVRAKWYPEVRHHCPNTPIILVGTKLDLR   120
          ************************************************************

hRac1     DDKDTIEKLKEKKLTPITYPQGLAMAKEIGAVKYLECSALTQRGLKTVFDEAIRAVLCPP   180
mRac1     DDKDTIEKLKEKKLTPITYPQGLAMAKEIGAVKYLECSALTQRGLKTVFDEAIRAVLCPP   180
zRac1a    DDKDTIEKLKEKKLTPITYPQGLAMAKEIGAVKYLECSALTQRGLKTVFDEAIRAVLCPP   180
          ************************************************************

hRac1     PVKKRKRKCLLL  192
mRac1     PVKKRKRKCLLL  192
zRac1a    PVKRRRRRCLLL  192
          ***:*:*:****
```

Rac1b: Protein similarity: 98%

```
hRac1     MQAIKCVVVGDGAVGKTCLLISYTTNAFPGEYIPTVFDNYSANVMVDGKPVNLGLWDTAG   60
mRac1     MQAIKCVVVGDGAVGKTCLLISYTTNAFPGEYIPTVFDNYSANVMVDGKPVNLGLWDTAG   60
zRac1b    MQAIKCVVVGDGAVGKTCLLISYTTNAFPGEYIPTVFDNYSANVMVDGKPVNLGLWDTAG   60
          ************************************************************

hRac1     QEDYDRLRPLSYPQTDVFLICFSLVSPASFENVRAKWYPEVRHHCPNTPIILVGTKLDLR   120
mRac1     QEDYDRLRPLSYPQTDVFLICFSLVSPASFENVRAKWYPEVRHHCPNTPIILVGTKLDLR   120
zRac1b    QEDYDRLRPLSYPQTDVFLICFSLVSPASFENVRAKWYPEVRHHCQTTPIILVGTKLDLR   120
          ********************************************* .*************

hRac1     DDKDTIEKLKEKKLTPITYPQGLAMAKEIGAVKYLECSALTQRGLKTVFDEAIRAVLCPP   180
mRac1     DDKDTIEKLKEKKLTPITYPQGLAMAKEIGAVKYLECSALTQRGLKTVFDEAIRAVLCPP   180
zRac1b    DDKDTIEKLKEKKLTPITYPQGLAMAKEIGAVKYLECSALTQRGLKTVFDEAIRAVLCPP   180
          ************************************************************

hRac1     PVKKRKRKCLLL  192
mRac1     PVKKRKRKCLLL  192
zRac1b    PVKKRKRKCSLL  192
          ********* **
```

B

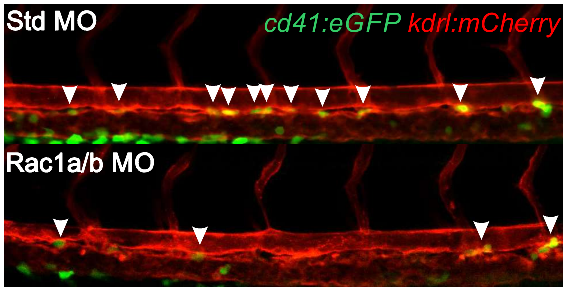


C

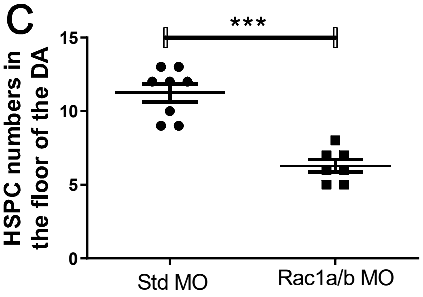

Sup 6

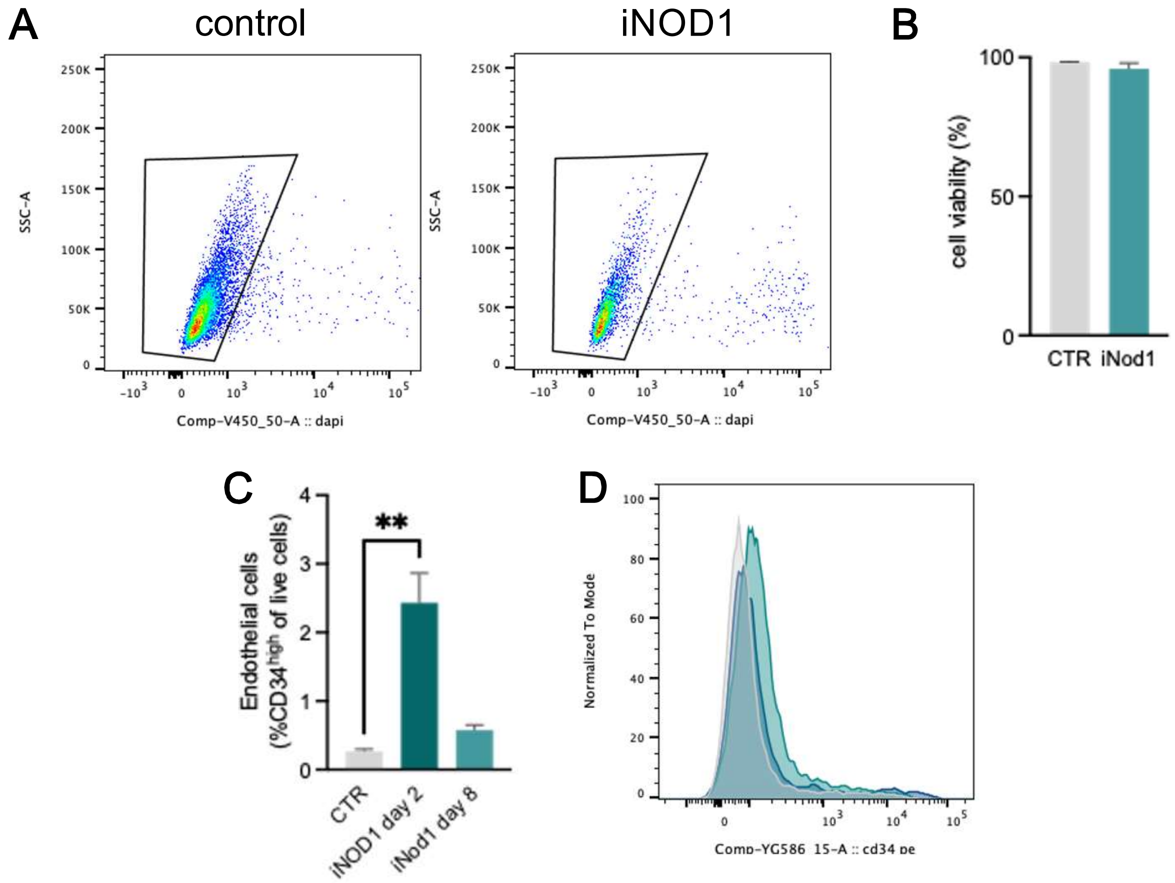
A
control
iNOD1
SSC-A
250K
200K
150K
100K
50K
0
$-10^3$
0
$10^3$
$10^4$
$10^5$
Comp-V450_50-A :: dapi
B
cell viability (%)
100
50
0
CTR
iNod1
C
Endothelial cells
(%CD34high of live cells)
4
3
2
1
0
**
CTR
iNOD1 day 2
iNod1 day 8
D
Normalized To Mode
100
80
60
40
20
0
Comp-YG586_15-A :: cd34 pe
