## Supplementary material for "Nod1-dependent NF-kB activation initiates hematopoietic stem cell specification in response to small Rho GTPases": Figure legends

**Figure1. NF-kB activation and NLRs expression during HE induction**

(A) HSPC developmental trajectory in zebrafish embryos.

(B) 16hpf *NF-kB: eGFP; kdrl:mCherry* double-transgenic embryos. White arrowheads denote *NF-kB*$^+$ regions in HE. Pictures were taken with 10x magnification.

(C) 20hpf *NF-kB:d2eGFP; kdrl:mCherry* double-transgenic embryos. Images were taken with 20x magnification.

(D) *kdrl:mCherry*$^+$ HE cells were purified by FACS at 22hpf for qPCR. Levels of indicated transcripts along x axis are shown relative to the housekeeping gene *ef1a* x 10000.

(E-G) UMAP depicting human endothelial scRNAseq dataset. aHEC = Arterial Hemogenic Endothelial cell; vEC = Venous Endothelial cell; aEC = Arterial Endothelial cell. F = *NOD1*; G = *CIITA*; H = *NLRX1*; I = *RIPK2*; J = *NFKBIA.*

**Figure 2. The non-inflammasome-forming NLR Nod1 is required for HSPC generation *in vivo***

(A) 16hpf *cd41:eGFP; kdrl:mCherry* double-transgenic embryos treated with Nodinitib-1 15 or 30µM and imaged at 48hpf. White arrowheads denote *cd41*$^+$*, kdrl*$^+$ HSPCs along the DA. Images were taken with 10X magnification.

(B) Quantification of *cd41*$^+$*, kdrl*$^+$ HSPCs from (A). Each data point represents total HSPCs per embryo.

(C) 48hpf *cd41:eGFP; kdrl:mCherry* double-transgenic embryos injected with Std MO or Nod1 MO1.

(D) Quantification of *cd41*$^+$*, kdrl*$^+$ HSPCs from (C).

(E) 48hpf *cd41:eGFP; kdrl:mCherry* double-transgenic embryos injected with Cas9 mRNA(control) or Cas9 mRNA with *nod1* gRNA.

(F) Quantification of *cd41*$^+$*, kdrl*$^+$ HSPCs from (E).

(G) *nod1*$^{+/+}$ and *nod1*$^{-/-}$ embryos were examined by WISH for *runx1* and *cmyb* expression in the aortic floor at 28hpf and 40 hpf. White arrowheads denote *runx1*$^+$ or *cmyb*$^+$ HSPCs.

(H) Quantification of *runx1*$^+$ or *cmyb*$^+$ HSPCs from (G). *p<0.05; **p<0.01 and ***p<0.001.

**Figure3. Nod1 programs the endothelium to become hemogenic**

(A) Schematic of Nodinitib-1 treatment at 16-24hpf or 24-48hpf.

(B) 16hpf or 24hpf *cd41:eGFP; kdrl:mCherry* double-transgenic embryos treated with 15uM Nodinitib-1 and imaged at 48hpf. Arrowheads denote *cd41*$^+$*, kdrl*$^+$ HSPCs along the DA. Images were taken with 10X magnification.

(C) Quantification of *cd41*$^+$*, kdrl*$^+$ HSPCs from (B). Each data point represents total HSPCs per embryo.

(D) WISH for *runx1* at 30hpf in uninjected embryos, or embryos injected with IEDAP 250ng/ul.

(E) Quantification of *runx1*$^+$ HSPCs from (D).

(F) WISH for *cmyb* at 42hpf in uninjected embryos, and embryos injected with IEDAP 250ng/ul or 400ng/ul.

(G) Quantification of *cmyb*$^+$ HSPCs from (F). ns = non-significant; **p<0.01 and ***p<0.001.

**Figure4. Ripk2 hyperactivation rescues Nod1-deficient HSPC phenotype**

(A) 48hpf *cd41:eGFP; kdrl:mCherry* double-transgenic embryos injected with Std MO or Ripk2 MO. Images were taken with 10X magnification.

(B) Quantification of *cd41*$^+$*, kdrl*$^+$ HSPCs from (A).

(C) *wildtype* and *ripk2*$^{-/-}$ embryos were examined by WISH for *runx1* or *cmyb* expression at 30hpf and 42hpf. White arrowheads denote *runx1*$^{+}$ or *cmyb*$^{+}$ HSPCs.

(D) Quantification of *runx1*$^{+}$ or *cmyb*$^{+}$ HSPCs from (C). Each dot represents total *runx1*$^{+}$ or *cmyb*$^{+}$ HSPCs per embryo.

(E) WISH for *runx1* at 30hpf in Control and ripk2$^{104ASP}$ embryos injected with Std MO or Nod1 MO. White arrowheads denote *runx1*$^{+}$ HSPCs.

(F) Quantification of *runx1*$^{+}$ HSPCs from (E). Each data point represents total *runx1*$^{+}$ HSPCs per embryo. $^{**}p<0.01$ and $^{***}p<0.001$.

(G) 5dpf *rag2:eGFP* embryos injected with Std MO, Nod1 MO2 or Ripk2 MO. Images were taken with 8X magnification. White arrowheads denote zebrafish thymus.

**Figure5. RNA Seq Transcriptomic analysis of the HE identify deregulated hematopoietic and inflammatory program**

(A) Schematic of experimental design. *Tg*(*kdrl:mCherry)* embryos were injected with or without Nod1 MO2. *kdrl:mCherry*$^{+}$ HE cells were purified from two conditions by FACS at 22hpf for RNA-seq.

(C) Enriched GO processes for significantly downregulated genes in Nod1 morphants versus wildtype embryos are listed in the tables.

(D) RNA-seq analysis from Nod1 morphants versus wildtype HE cells at 22hpf reveals 797 downregulated and 782 upregulated genes in Nod1-MO vs Std MO control.

(E) Heatmap displaying the extent of differential gene expression between 22hpf Nod1 morphants versus wildtype embryos. The genes incorporated in the heatmap represent all differentially expressed genes that belong to the GO term ‘immune system process’ downregulated genes.

(F) The genes incorporated in the heatmap represent all differentially expressed genes that belong to the GO term ‘embryonic hemopoiesis’ downregulated genes.

(G) The genes incorporated in the heatmap represent all differentially expressed genes that belong to the GO term ‘erythrocyte differentiation’ downregulated genes.

(H) The genes incorporated in the heatmap represent all differentially expressed genes that belong to the GO term ‘myeloid differentiation’ downregulated genes.

**Figure6. Nod1/Ripk2 signaling activates NF-kB within HE**

(A) 22hpf *NF-kB: eGFP; kdrl:mCherry* double-transgenic

embryos injected with Std MO or Nod1-MO2. Images were taken with 20x magnification.

(B) Flow cytometry analysis of *$GFP^+$/$mCherry^+$* expression in 22hpf *NF-kB:eGFP; kdrl:mCherry* double-transgenic embryos injected with Std-MO or Nod1-MO2.

(C) Conserved Nod1 Ser-177/181 (red) in human (hNOD1) and zebrafish (zNod1). Fluorescence microscopy imaging of *NF-kB:eGFP* embryos injected with mutated *ikkb (caIkkb)* versus *wt ikkb* mRNA control.

(D) WISH for *cmyb* at 40hpf in embryos injected with *ikkb* mRNA or $ikkb_{CA}$ RNA in addition to Std MO and Ripk2 MO. White arrowheads denote *$cmyb^+$* HSPCs.

(E) Quantification of *$cmyb^+$* HSPCs from (D). Each data point represents total *$cmyb^+$* HSPCs per embryo. *p<0.05 and ***p<0.001.

**Figure7. Rac1 activate the Nod1-Ripk2-NF-Kb axis *in vivo* to specify HSPCs**

(A) HE cells (*$flk^+$:mCherry*) from embryos at 22hpf were collected by FACS, and qPCR was performed for *rac1a, rac1b, rho* and *cdc42.* Levels of gene transcripts along the x-axis are shown relative to the housekeeping gene *ef1a* x 10000.

(B) 16hpf *cd41:eGFP; kdrl:mCherry* double-transgenic embryos treated with hydrochloride, rho kinase inhibitor III or ML141 and imaged at 48hpf. Arrowheads denote *cd41*$^{+}$, *kdrl*$^{+}$ HSPCs along the DA. All images were taken with 10X magnification.

(C) *cd41:eGFP; kdrl:mCherry* double-transgenic embryos were injected at 1 cell stage with *ikkb* or $ikkb_{CA}$ mRNA, chemically inhibited at 16hpf with hydrochloride, rho kinase inhibitor III or ML141 and quantified at 48hpf for *cd41*$^{+}$, *kdrl*$^{+}$ HSPCs. Each data point represents total HSPCs per embryo.

(D) WISH for *runx1* at 30hpf in Control and *ripk2104*$^{Asp}$ embryos injected with Cas9 only (control) or rac1a/b gRNA + Cas9. White arrowheads denote *runx1*$^{+}$ HSPCs.

(E) Quantification of *runx1*$^{+}$ HSPCs from(D). Each dot represents total *runx1*$^{+}$ HSPCs per embryo.

(F) WISH for *cmyb* at 42hpf in Control and ripk2104$^{Asp}$ embryos injected with Cas9 only (control) or rac1a/b gRNA + Cas9. White arrowheads denote *cmyb*$^{+}$ HSPCs.

(G) Quantification of *cmyb*$^{+}$ HSPCs from(F). Each data point represents total *cmyb*$^{+}$ HSPCs per embryo. ns= non-significant; *p<0.05; **p<0.01 and ***p<0.001.

**Figure8. Nod1 inactivation impairsthe formation of definitive human PSC-derived HSPCs**

(A) MegK = Megakaryocyte progenitor; Ery = Erythroid progenitor; HSPC = Hematopoietic Stem and Progenitor cells; Mono = Monocytes progenitor; Mono-Macs = Monocytes - Macrophages progenitor.

(B) Schematic of INod1 treatment from day2-day15 or day8-day15 on iPSCs. (C) Alive progenitors of CTR, iNod1 treated cells from day2 and iNod1 treated cells from day 8 at day 15.

(D) Quantification of live suspension cells from (B).

(E) Graphical abstract of the working model of this research. During HE induction, intracellular GTP-Rac1 activates Nod1 which activates Ripk2 upon binding through their CARD domains. Activated Ripk2 activates NF-kB subunits to translocate into the nucleus to start transcription. This process is essential for HSPC specification. LRRs = Leucine rich repeats; NBD = Nucleotide-binding domain; CARD = Caspase recruitment domain and KD = Kinase domain.
