## Supplementary material for "Nod1-dependent NF-kB activation initiates hematopoietic stem cell specification in response to small Rho GTPases": Material and Methods

**Table S1. Primer Sequences Used in This Study, Related to Experimental Procedures**

| Gene | Accession | Name | Nucleotide sequence (5'-3') | Use |
|---|---|---|---|---|
| *nod1* | XM_002665060.6 | F | GGCACGAACAATTTCGTTTT | gRNA validation |
| | | R | CTGTTTTAGTGCTGCTGCGC | |
| | | F | GCGGTATTGAGGTTCTGGCT | qPCR |
| | | R | TGTGGTTTTGGTAAAGGCCCA | |
| | | F | TTAGATGAAGCGGTGTGCTG | Sa12613 mutant line PCR |
| | | R | TGCAGATATGCCGTTAGCTG | |
| | | F | GTGAAGGTGTTGGGGTGAGT | Sa17969 mutant line PCR |
| | | R | CAAACAAGTGACCACCATGC | |
| | | F | GATCTTTCATTTCATTTTTCAGAGCCCGA | Morpholino validation |
| | | R | GATCTCGGGCTCTGAAAAATGAAATGAAA | |
| *ripk2* | NM_194411.2 | F | GGGTCTGCCGTCATCATTAAT | Z40 mutant line validation |
| | | R | GTGAGGGGTTGTATGGCAAGA | |
| | | F | GGGTCTAGTACGTAGGCTGGA | qPCR |
| | | R | CACCGGTAATGTGCTGGTGA | |
| *rac1a* | NM_199771.1 | F | CCGCTCTTGTTTTGCGTGTT | qPCR |
| | | R | TTTTTCCCACAGCCCCGTCC | |
| | | F | ACTCATGGATATCGGCAAGC | gRNA validation |
| | | R | CGGTCGAAGCCTGTCATAAT | |
| *rac1b* | NM_001039818.1 | F | GGGGTTTTTCATCAGTTCCGC | qPCR |
| | | R | TTTTACCCACAGCCCCGTCA | |
| | | F | AGGAAGCTGCCATGGTGTTA | gRNA validation |
| | | R | CATGTCTGCAGGTTTGTGCT | |
| *nod2* | NM_001328044.1 | F | ACACA CC CACAACAGGTTC | qPCR |
| | | R | GAAGAGGGACTGCGATGCAA | |
| *cdc42* | NM_001018120.2 | F | CTCTGACGCAGCGAGGTC | qPCR |
| | | R | TGCGTTTCAGGAGGTTCGAG | |
| *rho* | NM_131084.1 | F | CCGGAGCCCATACGAATACC | qPCR |
| | | R | AGGAAGAACATGTAGGCCGC | |
| *ef1a* | NM_131263.1 | F | GAGAAGTTCGAGAAGGAAGC | qPCR |
| | | R | CGTAGTATTTGCTGGTCTCG | |
| *ciita* | XM_005163915.4 | F | GCACTGTGGTTCAGACAGGA | qPCR |
| | | R | CAACCGTACCATCAGCAGGT | |
| *nlrc5* | NM_001386270.1 | F | TCCTTCCTGTCATGTTGTCTC | qPCR |
| | | R | TCAGCTTGGTGCCTGAGTTC | |
| *nlrx1* | XM_680389.9 | F | TCCACACAGTGCATCTGTACC | qPCR |

| | | | |
|---|---|---|---|
| | | R | CAGAGATGTCCGAACCCTCG |

**Table S2. Morpholinos Used in This Study, Related to Experimental Procedures**

| Name | **Sequence (5'-3')** | **Concentration injected** | **Reference** |
|---|---|---|---|
| Std-MO | CCTCTTACCTCAGTTACAATTTATA | 0.4mM | - |
| Nod1-MO1 | TTTCATTTCATTTTTCAGAGCCCGA | 0.8mM | This study |
| Nod1-MO2 | ACCAAATAAACATTACCTGGTCTGT | 1mM | (Oehlers et al., 2011) |
| Ripk2-MO | GCTCCATGTTTCTGGACATTAGGAG | 1mM | This study |
| Nod2-MO | GTTTAAGGTGGTATTACCTGTTGTG | 1.24mM | (Oehlers et al., 2011) |
| Rac1a-MO | CCACACACTTTATGGCCTGCATCTG | 0.28mM | (Mikdache et al., 2020) |
| Rac1b-MO | CCACACACTTGATGGCCTGCATGAC | 0.28mM | (Epting et al., 2015) |

**Table S3. Chemicals Used in This Study, Related to Experimental Procedures**

| **Name** | **Use** | **Concentration used** | **Reference** |
|---|---|---|---|
| C12-iE-DAP | Nod1 agonist | 250ng/ul – 400ng/ul | InvivoGen (tlrl-c12dap) |
| Rho Kinase Inhibitor III | Rho inhibitor | 15.14uM | (Weiser et al., 2009) |
| Hydrochloride | Rac1 inhibitor | 48.96uM | (Nussbaum et al., 2013) |
| ML141 | Cdc42 inhibitor | 38.66uM | (Stanganello et al., 2015) |

### Zebrafish husbandry and strains

Zebrafish embryos and adults were mated, staged, raised, and processed as described (Westerfield, 2000) in a circulating aquarium system maintained at 28$^{o}$C. Two *nod1* mutant zebrafish strains (*sa12613*, and *sa17969*) were obtained from the Zebrafish International Resource Center. *ripk2*$^{-/-}$ zebrafish mutants were kindly donated by Michael Jurynec (Jurynec et al., 2018). Other zebrafish lines used in this study were: WT *AB**, transgenic *Tg(cmyb:eGFP)*$^{zf169}$, *Tg(kdrl:HsHRAS-mCherry)*$^{s896}$ (referred to as *kdrl:mCherry* throughout the manuscript) (Bertrand et al., 2010), *Tg (-6.0itga2b:eGFP)*$^{la2}$ (referred to as *cd41:eGFP* throughout the manuscript) (Lin et al., 2005), *Tg(NFkB:eGFP)*$^{nc1}$ (Kanther et al., 2011), *Tg(Rag2:eGFP)*$^{zdf8}$ (Langenau et al., 2003) and various intercrosses of these lines were utilized.

**FACS isolation of zebrafish embryonic cell populations**

FACS was performed as previously described (Barakat et al., 2022). To isolate 2hpf, 4hpf, 24hpf, 48hpf embryo cells for qPCR analysis, approximately 150 2hpf, 4hpf, 24hpf and 48hpf *AB* wildtype zebrafish embryos were dechorionated with pronase, anesthetized in 1% tricaine, gently shaken at 28$^{o}$C for 5 or 10 mins with 0.05mg/ml liberase TM (Roche) in PBS solution with $Ca2^{+}$ and $Mg2^{+}$. The resulting cell suspension was filtered and collected.

To isolate endothelial cells at 24hpf for qPCR analysis, approximately 150 24hpf *kdrl:mCherry* zebrafish embryos were dechorionated with pronase, anesthetized in 1% tricane, gently shaken at 28$^{o}$C for 5 mins with 0.05mg/ml liberase TM (Roche) in PBS solution with $Ca2^{+}$ and $Mg2^{+}$. The resulting cell suspension was filtered and stained with Sytox Red to exclude dead cells. *kdrl*+ cells were sorted with BD FACS Aria III.

To isolate whole embryo cells at 24hpf or 48hpf for qPCR analysis, approximately 150 24hpf and 48hpf *AB* wildtype zebrafish embryos were dechorionated with pronase, anesthetized in 1% tricane, gently shaken at 28$^{o}$C for 5 or 10 mins with 0.05mg/ml liberase TM(Roche) in PBS solution with $Ca2^{+}$ and $Mg2^{+}$. The resulting cell suspension was filtered and collected.

To isolate muscle cells at 24hpf and 48hpf for qPCR analysis, approximately 150 24hpf and 48hpf *phldb:eGFP* zebrafish embryos were dechorionated with pronase, anesthetized in 1% tricane, gently shaken at 28$^{o}$C for 5 or 10 mins with 0.05mg/ml liberase TM(Roche) in PBS solution with $Ca2^{+}$ and $Mg2^{+}$. The resulting cell suspension was filtered and stained with Sytox Red to exclude dead cells. *phldb*$^{+}$ cells were sorted and collected with BD FACS Aria III.

To isolate myeloid cells at 24hpf and 48hpf for qPCR analysis, approximately 150 24hpf and 48hpf *mpeg:eGFP* and *mpx:eGFP* zebrafish embryos were dechorionated with pronase, anesthetized in 1% tricane, gently shaken at 28$^{o}$C for 5 or 10 mins with 0.05mg/ml liberase TM(Roche) in PBS solution with $Ca2^{+}$ and $Mg2^{+}$. The resulting cell suspension was filtered and stained with Sytox Red to exclude dead cells. *mpeg*$^{+}$ (macrophages) or *mpx*$^{+}$ (neutrophils) cells were sorted and collected with BD FACS Aria III.

To isolate HSPCs, endothelial cells, hematopoietic cells at 48hpf for qPCR analysis, approximately 150 48hpf *kdrl:mCherry; cd41:eGFP* double transgenic zebrafish embryos were dechorionated with pronase, anesthetized in 1% tricane, gently shaken at 28$^{o}$C for 10mins with 0.05mg/ml liberase TM (Roche) in PBS solution with $Ca2^{+}$ and $Mg2^{+}$. The resulting cell suspension was filtered and stained with Sytox Red to exclude dead cells. *cd41*$^{+}$*,kdrl*$^{+}$(HSPCs); *cd41*$^{-}$*:kdrl*$^{+}$(endothelial cells); *cd41*$^{+}$*,kdrl*$^{-}$(hematopoietic cells) cells were sorted and collected with BD FACS Aria III.

**Quantitative RT-PCR Analysis**

RNA was isolated from FACS sorted cells with RNeasy Micro Kit (Qiagen), and cDNA was generated with qScript Supermix (Quanta BioSciences) or iScript gDNA Clear cDNA

Synthesis Kit (BioRad). Primers to detect zebrafish transcripts are described in **Table S1**. qPCR was performed with CFX Connect Real-Time System (BioRad).

**Morpholino injection**

MOs used in this study are described in **Table S2.** MOs were resuspended in nuclease-free water at 2mM, diluted with nuclease-free water and 1ul phenol red solution. 1nl was injected in the yolk ball of one-cell-stage embryos using PLI-90A pico-injector warner instruments.

**gRNA design and injection**

To identify CRISPR gRNA sites in *nod1, rac1a* and *rac1b*, targeted genomic and coding sequences were found on <emsemble.org>. In the targeted sequence, PAM sequence was located in 5' exons. The cutting site was predicted to be 3bp into the target sequence upstream from the PAM. 20bp upstream of PAM was identified as gRNA and ordered from Synthego. (*nod1* gRNA: GACUGUUCACAGAGAGCUGC; *rac1a* gRNA: ACCAGUAAACCUGGGAUUGU; *rac1b* gRNA: CUGCGAAUGUGAUGGUGGAU). gRNA efficiency was then determined by injection of 25pg gRNA along with 300pg Cas9 mRNA into the first cell of developing embryos. Whole embryos were hot shot at 48hpf for genomic DNA. Targeted sequences were then PCR amplified using primers described in **Table S1**. Amplicons were Sanger sequenced and gRNA efficiency was then analyzed using Synthego ICE Analysis. Following high efficiency gRNA knockout results, 25pgRNA was injected with 300pg Cas9mRNA into the first cell of developing embryos for various experiments in this research.

**In Situ Hybridization**

WISH was performed as described (Thisse et al., 1993). Probes used for *cmyb*, *runx1, efnb2a and kdrl* transcripts were generated using the DIG RNA Labeling Kit (Roche Applied Science) from linearized plasmids. Embryos were imaged using a Leica M165FC stereomicroscope equipped with a DFC295 color digital camera (Leica) and FireCam software (Leica).

**Enumeration of HSPCs**

Zebrafish embryos were performed with WISH for *runx1* and *cmyb* at noted stages. Positive cells are imaged and manually counted. Confocal microscopy was performed on *cd41:eGFP; kdrl:mCherry* double-transgenic embryos. Z sections of the DA region were imaged on a Ziess LSM700 Laser Scanning Confocal and double positive cells were manually counted.

**NF-kB reporter quantification analysis**

To quantify the *NFkB* expression in endothelial cells from Standard-MO injected and Nod1-MO2 injected *AB* wildtype embryos at 22hpf, confocal microscopy was performed on *NFkB:eGFP;kdrl:mCherry* double transgenic control embryos or nod1 morphants. Z sections of the DA region were imaged on a Zeiss LSM700 Laser Scanning Confocal. *kdrl:mCherry* positive DA regions were selected on the images and analyzed for green intensity using color histogram in ImageJ.

**Flow cytometry**

To quantify the *NFkB* expression in endothelial cells from Standard-MO injected and Nod1-MO2 injected *AB* wildtype embryos at 22hpf, approximately 150 23hpf *NFkB:eGFP;kdrl:mCherry* double transgenic control embryos or nod1 morphants were dechorionated with pronase, anesthetized in 1% tricaine, gently shaken at 28$^{o}$C for 5 mins with 0.05mg/ml liberase TM (Roche) in PBS solution with $Ca2^{+}$ and $Mg2^{+}$. The resulting cell suspension was filtered and stained with Sytox Red to exclude dead cells. Flow cytometric acquisitions were performed on a Melody (BD) and analyses were performed using FlowJo software (v10.3, Tree Star).

**RNA sequencing (RNA-seq) Preparation**

*AB* wildtype x *kdrl:mCherry* zebrafish embryos were injected with Standard-MO, Nod1-MO2, Nod2-MO and Ripk2-MO at one-cell stage. Approximately 150 per morpholino injection 24hpf *kdrl* $^{+}$ zebrafish embryos were screened with Leica M165FC stereomicroscope, dechorionated with pronase, anesthetized in 1% tricane, gently shaken at 28$^{o}$C for 5 mins with 0.05mg/ml liberase TM(Roche) in PBS solution with $Ca2^{+}$ and $Mg2^{+}$. The resulting cell suspension was filtered and stained with Sytox Red to exclude dead cells. *kdrl*$^{+}$ cells were sorted and collected with BD FACS Aria III. RNA was then isolated and purified with RNeasy Micro Kit (Qiagen). Total RNA was assessed for quality and quantity using an Agilent 2100 Bioanalyzer with RNA 6000 Pico Kit. Samples were then used to generate RNA sequencing libraries using NEBNext Single Cell/Low Input RNA Library Prep Kit (NEB #E6420S/L) for Illumina. Triplicates RNA samples of each condition were processed following manufacturer's instructions.

**Pluripotent Stem Cells maintenance**

hPSCs were cultured in StemPro hESC SFM (Gibco) in presence of 20ng/ml bFGF (R&D) on Vitronectin (ThermoFisher Scientific) coated plates. hPSCs cells were passaged using ReLeSR (Stem Cell Technologies). Media change was performed every day and cells passaged every 3–4 days at a ratio of 1:6-1:10.

**Pluripotent Stem Cells differentiation to hematopoietic progenitors**

hPSCs were differentiated with a modified version of our previously published protocol (Fidanza *et al.*, 2020). hPSCs were made into cluster by using ReLeSR as for passaging and resuspended in Day 0 differentiation medium, containing 10 ng/ml. Cell clusters were cultured onto Cell Repellent 6 wells Plates (Greniner) to support embryoid bodies (EBs) formation. At day 2 the media was changes and 3 μM CHIR (StemMacs) was added; to some of the wells Nod1 inhibition was started by addition of Nodinitib-1 (Cayman Chemicals). At day 3, EBs were transferred into fresh media supplemented with 5 ng/ml bFGF and 15 ng/ml VEGF. At day 6 media was changed for final haematopoietic induction in SFD medium supplemented with 5 ng/ml bFGF, 15 ng/ml VEGF, 30 ng/ml IL3, 10 ng/ml IL6, 5 ng/ml IL11, 50 ng/ml SCF, 2 U/ml EPO, 30 ng/ml TPO, 10 ng/ml FLT3L and 25 ng/ml IGF1.

**CD34 isolation**

At day 8 CD34+ cells were isolated with CD34 Magnetic Microbeads (Miltenyi Biotec) according to their manufacturing protocol. Briefly, EBs were dissociated using Accutase (Life Techonologies) at 37°C for 30'. Cells were centrifuged and resuspended in PBS + 0.5% BSA + 2mM EDTA in presence of Fcr blocker and magnetic anti-CD34 and incubated at 4°C for 30'. CD34+ cells were isolated using MS column with the aid of a magnet. After centrifugation, cells were resuspended in SFD media, counted, and plated for OP9 coculture.

**OP9 coculture and flow cytometry**

OP9 cells were cultured in α-MEM, sodium bicarbonate (Gibco) and 20% serum (Gibco); they were passaged with Trypsin every 3-4 days. The day preceding the co-culture,

45.000 OP9 cells were plated in each 12 well plates' well in SFD media. The day of the co-culture 20.000 CD34+ cells were plated in each well and culture in SFD media supplemented with 5 ng/ml bFGF, 15 ng/ml VEGF, 30 ng/ml IL3, 10 ng/ml IL6, 5 ng/ml IL11, 50 ng/ml SCF, 2 U/ml EPO, 30 ng/ml TPO, 10 ng/ml FLT3L and 25 ng/ml IGF1. Cytokines were topped up once during one week of coculture. After 7 days of co-culture, human hematopoietic suspension cells were collected by gently flashing the coculture and aspirating the whole media and counted. For flow cytometry, adherent cells were detached using Accutase and added to the suspension cells, after which they were centrifuged and resuspended in PBS + 0.5% BSA + 2mM EDTA and stained with antibodies for 30' at room temperature gently shaking using CD34-Pe (1:200, 4H11, Invitrogen) and CD43-APC (1:100, eBio84-301, Invitrogen); DAPI was used for live-dead gating. Flow cytometry data were collected using DIVA software (BD) and analysed using FlowJo 10.8.1 software (BD).

**Single Cell RNAseq analysis**

Gene expression of target genes was analysed in human iPSCs derived cells (Fidanza et al., 2020) and in human AGM collected *in vivo* from Carnegie stage CS12-14 (Zeng et al., 2019). For the human iPSCs derived dataset, gene expression values were obtained using the web portal at https://lab.antonellafidanza.com/, for the *in vivo* dataset the data were analyzed using Seurat R package (Stuart et al., 2019).

**Statistical Analyses**

Data were analyzed by unpaired T-test, or two-way ANOVA with Tukey's post-test in Graphpad Prism 5. In all figures, middle black bars denote the mean and error bars represent S.E.M. *p<0.05, **<0.01, ***p<0.001, ns=not significant. ND=not detected.
